# Fast rate coding in heterogeneous spiking networks

**DOI:** 10.64898/2026.10.30.763048

**Authors:** Kasper Smeets, Shuqi Wang, Valentin Schmutz, Wulfram Gerstner

**Affiliations:** School of Life Sciences and School of Computer and Communication Sciences, EPFL, Lausanne, Switzerland; Department of Mathematics, ETH Zürich, Zürich, Switzerland

## Abstract

A network of spiking neurons is said to operate in the rate coding regime if its dynamics can be accurately described by a continuous rate network. But how can discrete spike-based communication between neurons give rise to continuous rate-based dynamics? Computational studies showing that heterogeneous spiking networks can approximate the dynamics of recurrent neural networks (RNNs), a special case of rate network, assume that spikes are approximately Poisson or that network dynamics are slow. Here, we show that neither assumption is necessary: heterogeneous networks of non-Poisson neurons can produce fast rate-based dynamics. To demonstrate this, we show that signal transmission in feedforward networks of integrate-and-fire-type neurons can be described by a history-dependent rate network that is asymptotically exact as network size grows, even if heterogeneity persists, while linear-nonlinear rate networks fail. By providing a constructive model of fast rate coding in heterogeneous networks, this work lays the foundation for extending rate-based analyses of neural recordings beyond the assumption of slow dynamics or Poisson rate codes.

## Introduction

Neuronal responses are commonly analyzed in terms of instantaneous firing rates. These continuously valued variables are useful for inferring each neuron’s involvement over time during, e.g., sensory stimulation (Roitman and Shadlen, 2002), decision making (Luo et al., 2025), or generation of behavior (Churchland and Shenoy, 2024). While widely adopted, a critical question remains as to whether and when instantaneous firing rates — such as those obtained by averaging spike trains across trials or by combining population-level variables in single trials — can be representative of neuronal communication. This is the core question of rate coding (Brette, 2015). More generally, when and how can noisy, discrete spike-based communication be fully described by deterministic, continuous rate-based interactions without loss of function? We define rate coding in a spiking network by the possibility to construct a one-to-one mapping, one spiking neuron to one rate unit, from a spiking network to a rate network, such that the rate network reproduces the spiking network’s dynamics under identical external stimulation. The ability to construct such a mapping then supports the mechanistic interpretability of rate-based analyses of experimental data.

While a downstream neuron does not have access to the trial-averaged spike train of a presynaptic neuron, the trial-averaged signal has been argued to be present in a single trial if there exists a pool of many (nearly) identical presynaptic neurons that drive the postsynaptic neuron (Perkel and Bullock, 1968; Shadlen and Newsome, 1994). Indeed, theoretical mappings have shown that the pooled spiking activity of infinitely many neurons driven by a common stimulus can be described for many different spiking models through deterministic population density equations (Knight, 1972; Gerstner, 2000; Brunel, 2000; Pakdaman et al., 2010) or (next-generation) neural mass models (Montbrió et al., 2015; Pietras et al., 2019; Byrne et al., 2020), and that this population activity can also be understood as the trial-averaged spiking activity of a single neuron (Herrmann and Gerstner, 2001; Naud and Gerstner, 2012).

For networks without pools of similarly tuned neurons, existing theoretical mappings require either that spike trains can be seen as generated by linear-nonlinear Poisson processes (Schmutz et al., 2025) or that network dynamics are slow enough to be considered “quasi-stationary” (see e.g., (Dayan and Abbott, 2001, Chapter 7.2), (Gerstner and Kistler, 2002, Chapter 6.3), (Ermentrout and Terman, 2010, Chapter 11.1)). In both cases the resulting rate network is described by ordinary rate equations, such as those defining standard recurrent neural networks (RNNs). These theoretical mappings enable training methods available for RNNs to be applied to spiking networks satisfying these assumptions (DePasquale et al., 2016; Abbott et al., 2016; Kim and Chow, 2018; DePasquale et al., 2023; Newton and Nicola, 2025), and explain how RNNs can sometimes be directly transferred to spiking networks without loss of function (Kim et al., 2019; Cimeša et al., 2023). However, neither of the two assumptions is generally valid for biological networks, since neurons do not generally obey Poisson statistics (Reich et al., 1998; Keat et al., 2001; Simoncelli et al., 2004), and fast responses to incoming stimuli are commonly observed in the brain (Thorpe et al., 1996; Marálek et al., 1997; Fabian et al., 2023; Zeraati et al., 2023). Non-Poisson behavior particularly emerges when neurons are driven by fast time-varying signals (Mainen and Sejnowski, 1995), consistent with the theory of spiking neurons with refractoriness (Knight, 1972; Perkel et al., 1967; Gerstner, 2000; Brunel et al., 2001; Moreno-Bote and Parga, 2004; Tchumatchenko et al., 2011). The limitations of these assumptions are also highlighted by the finding that, outside of the quasi-stationary state, training methods assuming ordinary rate dynamics no longer work on spiking networks (Kim and Chow, 2018; Newton and Nicola, 2025). More generally, describing networks of spiking neurons by standard rate models may cause invalid interpretations of neuronal communication (Amarasingham et al., 2015; Chandrasekaran et al., 2018; Aghamohammadi et al., 2025).

In this paper, we show that an accurate rate description is possible for heterogeneous spiking networks even when network dynamics evolve on a time scale comparable to the membrane time constant and neuronal spike trains have highly non-Poisson characteristics. We study a network of noisy spiking neurons (Leaky Integrate-and-Fire (LIF) or Spike Response Model (SRM_0_) (Gerstner, 1995, 2000; Kass and Ventura, 2001)) with reset or refractoriness and construct a corresponding rate network to describe neuronal communication. Crucially, our rate network differs from standard rate models by including an explicit history dependence that is necessary to capture the rich short-term response properties shaped by the voltage reset and refractoriness following spike emission. In our rate network, each individual spiking neuron is replaced by a single history-dependent rate unit, which we visualize each as representing a *virtual population* — even though the number of rate units is identical to the number of spiking neurons in the original network. We find that replacing each *single* spiking neuron by a virtual population as a *single* rate unit yields an accurate description of the spiking network, whereas replacement with ordinary rate units, or linear-nonlinear generalizations thereof, does not. Overall, our results show that rapid time-dependent coding with a continuous rate variable is possible in model networks of spiking neurons, even if there are no pools of similar neurons and spike trains are not Poisson.

## Results

Showing that spike-based interactions in a spiking network can be reduced to rate-based interactions, i.e., demonstrating the causal role of firing rates (Brette, 2015), requires constructing a rate network with the same connectivity that reproduces the dynamics of the spiking network. Formally, starting from a spiking network of *N* spiking neurons, a successful mapping to a rate network requires three components. First, it requires a corresponding rate network of *N* rate units, with each unit *i* defined by a mathematical expression of how the instantaneous firing rate *r*_*i*_(*t*) depends on past firing rates *r*_*j*_(*t*^*′*^), for *t*^*′*^ *< t*, of presynaptic rate units *j* = 1, 2, …, *N* :

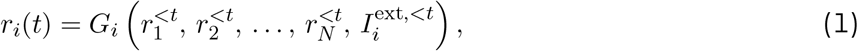

where 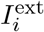 denotes external input received by neuron *i* and *G*_*i*_ is a function (or functional, dynamical system, etc.) of the past rates of the *N* units in the network. Second, a one-to-one transformation that tells us how parameters of a given spiking neuron *i* determine *G*_*i*_. Third, a theory that identifies assumptions under which the mapping is accurate and therefore meaningful, in the sense that the rate network produces (approximately) the same network dynamics as the spiking network when driven with the same external input.

Ordinary rate networks formulated with differential equations (Miller and Fumarola, 2012), as well as linear-nonlinear generalizations thereof (see e.g. Ostojic and Brunel (2011)), are special cases of Eq. (1); see Methods.

### From weighted sums of spikes to weighted sums of rates

To derive an expression for *G*_*i*_, and to explain how noisy spike-based communication can give rise to deterministic rate-based dynamics, we consider a single postsynaptic neuron *i* embedded in a feedforward network (Fig. 1a) which receives inputs from *N* presynaptic spiking neurons (Fig. 1b, top). Each time 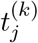 a presynaptic neuron *j* emits a spike, it induces a short postsynaptic current (through, e.g., AMPA or GABA channels) that affects the post-synaptic membrane potential. Formally, we model these pulses as 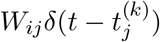, where *W*_*ij*_ is the synaptic weight from neuron *j* to neuron *i* and *δ* denotes the unit impulse function. The sequence of spikes emitted by a presynaptic neuron *j* at times 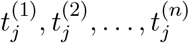 forms a spike train 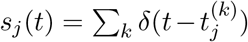. The spike trains of several presynaptic neurons are integrated by the postsynaptic neuron *i* according to

**Figure 1.**
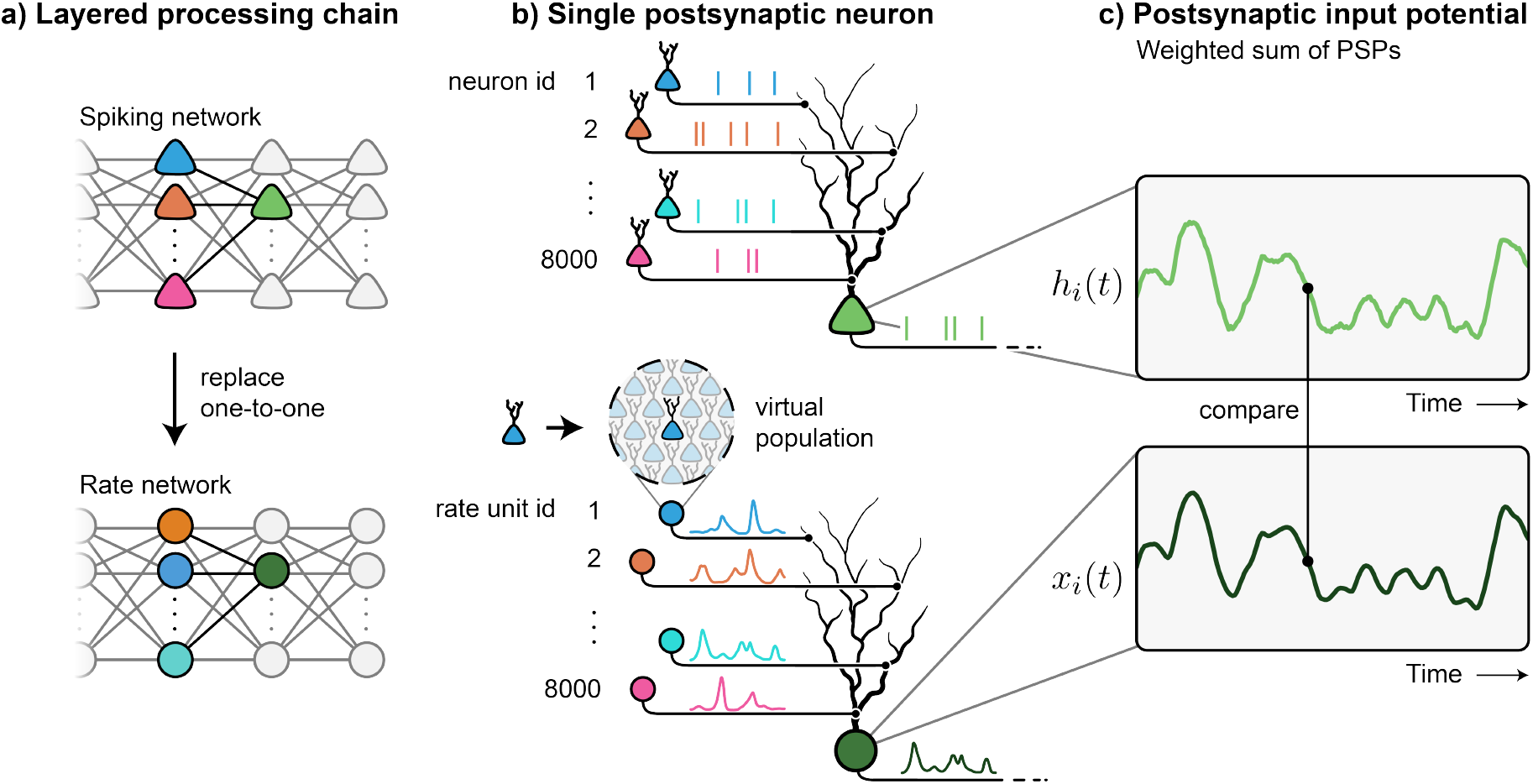
From spiking networks to networks of “virtual population” rate-units. **a)** One-toone, neuron-to-unit mapping from a heterogeneous network of spiking neurons (filled triangles, top) to a network of rate units (filled circles, bottom). Different colors indicate that neurons have different parameters. **b)** The green postsynaptic neuron receives input from many presynaptic neurons in the form of discrete spikes. To define our rate network, we imagine that each spiking neuron is replaced by a virtual population of identical neuronal copies. **c)** The postsynaptic potentials cause by spike arrivals are summed and yield a predictable spike-based input potential (top). In the rate network, the rate-based input potential of the green postsynaptic rate unit (bottom) is nearly identical to that of the green neuron in the spiking network. Simulation using 8000 spiking neurons (top) or 8000 rate units (bottom) as inputs to a single neuron.

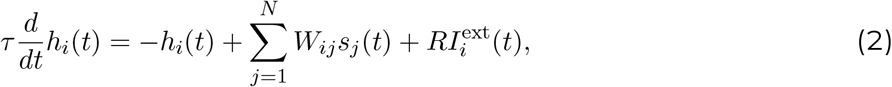

where *τ* is the membrane time constant, set to 10 ms, and *R* is the resistance of the membrane. We call the variable *h*_*i*_(*t*) the *input potential* of neuron *i* because it represents the contributions of the inputs to the membrane potential. That is, it represents the membrane potential without the action potentials and the spike after-potentials caused by the neuron’s own spikes, such as the voltage reset in integrate-and-fire-type neuron models (Fig. 1c, top).

The key insight for mapping a heterogeneous spiking network to a rate network follows from the observation that the input potential *h*_*i*_(*t*), given by Eq. (2), is a weighted sum of presynaptic spike trains *s*_1_(*t*), …, *s*_*N*_ (*t*) (plus the external input) after membrane filtering. If spike times are variable (as it is often observed in experiments (Shadlen and Newsome, 1994, 1998)) and if this variability is uncorrelated between neurons, the postsynaptic sum *h*_*i*_(*t*) can behave almost like a deterministic quantity because the summation runs over thousands of presynaptic inputs. The weighted sum of filtered spikes effectively averages out uncorrelated spiking variability, resulting in the input potential *h*_*i*_(*t*) behaving as if each presynaptic neuron *j* was communicating its underlying instantaneous firing rate (Dayan and Abbott, 2001). This statistical phenomenon is known as the concentration of measure phenomenon, and has previously been used to construct a mapping for networks of linear-nonlinear-Poisson spiking neurons (Schmutz et al., 2025).

The insight presented above suggests that to derive a rate network from a spiking network, we should consider a rate network that has the same synaptic weights as the spiking network (Fig. 1a, bottom) but replace each input potential *h*_*i*_(*t*), Eq. (2), by a rate-based counterpart that integrates the time-varying presynaptic firing rates *r*_1_(*t*), …, *r*_*N*_ (*t*) instead of the spike trains *s*_1_(*t*), …, *s*_*N*_ (*t*) (Fig. 1b, bottom). This rate-based input potential, denoted by *x*_*i*_(*t*), evolves according to

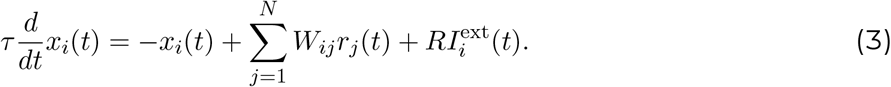

In a large spiking network, the spike-based input potentials *h*_*i*_(*t*), obeying Eq. (2), are expected to concentrate around their rate-based counterparts *x*_*i*_(*t*), obeying Eq. (3) (Fig. 1c).

In the spiking model of Eq. (2), spikes are generated stochastically and each spike causes a spike after-potential (or reset of the membrane potential). In the rate model of Eq. (3), no such reset occurs and communication happens with temporally smooth variables *r*_*i*_(*t*). To obtain an expression for the *G*_*i*_ in the rate model of Eq. (1), we first need to estimate the instantaneous firing rate of each spiking neuron. To do so, we imagine that we replace each spiking neuron *i* in the spiking network by a “virtual population” of infinitely many identical copies of neuron *i*, sharing the same neuronal parameters of neuron *i* and all receiving the same input (Fig. 1b, bottom). Each rate unit *i* in the rate network is then defined by the input-output relation of this virtual population. That is, *G*_*i*_ as in Eq. (1) is defined by the population-averaged spiking activity in response to the time-varying (rate-based) input potential *x*_*i*_(*t*).

In the case of renewal-type models of spiking neurons with escape noise (Gerstner, 1995), such as for LIF and SRM_0_, the virtual population approach leads to a *history-dependent rate network*, where each firing rate takes the shape 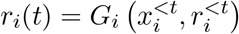, which explicitly depends on a unit’s own firing rate history *r*^*<t*^ (Methods), allowing it to capture refractory effects and fast transient responses (Gerstner, 2000). We emphasize that the input potential *x*_*i*_ summarizes the effect of the past firing rates of presynaptic neurons 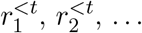 and external input 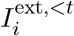 cf. Eq. (1). The history-dependent rate network naturally reduces to standard rate equations under quasi-stationary dynamics, which we call the *ordinary rate network*. In the ordinary rate network, each rate unit *i* is also governed by a rate-based input potential *x*_*i*_(*t*) (Eq. 3) but has a its firing rate equation given by *r*_*i*_(*t*) = *F*_*i*_ (*x*_*i*_(*t*)), with *F*_*i*_ being the ‘f-I curve’ of spiking neuron *i*, i.e., the function that gives the stationary firing frequency of a neuron as a function of the constant input it receives (Miller and Fumarola, 2012).

### History-dependent rate network captures fast, single-trial spiking network dynamics

To illustrate our mapping, we start with a simple spiking network consisting of *N* = 8000 SRM_0_ presynaptic spiking neurons projecting to a single postsynaptic output neuron (Fig. 2a, top). To construct the corresponding history-dependent rate network, the *N* = 8000 presynaptic neurons are replaced in a one-to-one manner by *N* = 8000 history-dependent rate units, which project to a single output unit (Fig. 2a, bottom). Both networks are driven by the same *P* = 20 independent step-like input signals, changing their amplitude every 50 ms. In the spiking network, the input potential *h*^out^(*t*) of the output neuron extracts one of the *P* signals from the presynaptic spikes through a weighted summation. Analogously, in the rate network, the input potential *x*^out^(*t*) of the output rate unit extracts the signal from the presynaptic rates. In addition to the heterogeneity of neuronal parameters, each neuron/unit has its own stimulus tuning, i.e., each neuron responds preferentially to one out of 2^20^ (more than one million) possible binary combinations of the 20 signals (Hopfield, 1982, 1984). The resulting neural responses are highly diverse (Fig. 2b-c) with spike trains that are only weakly correlated (Fig. 2f-g; long simulation).

**Figure 2.**
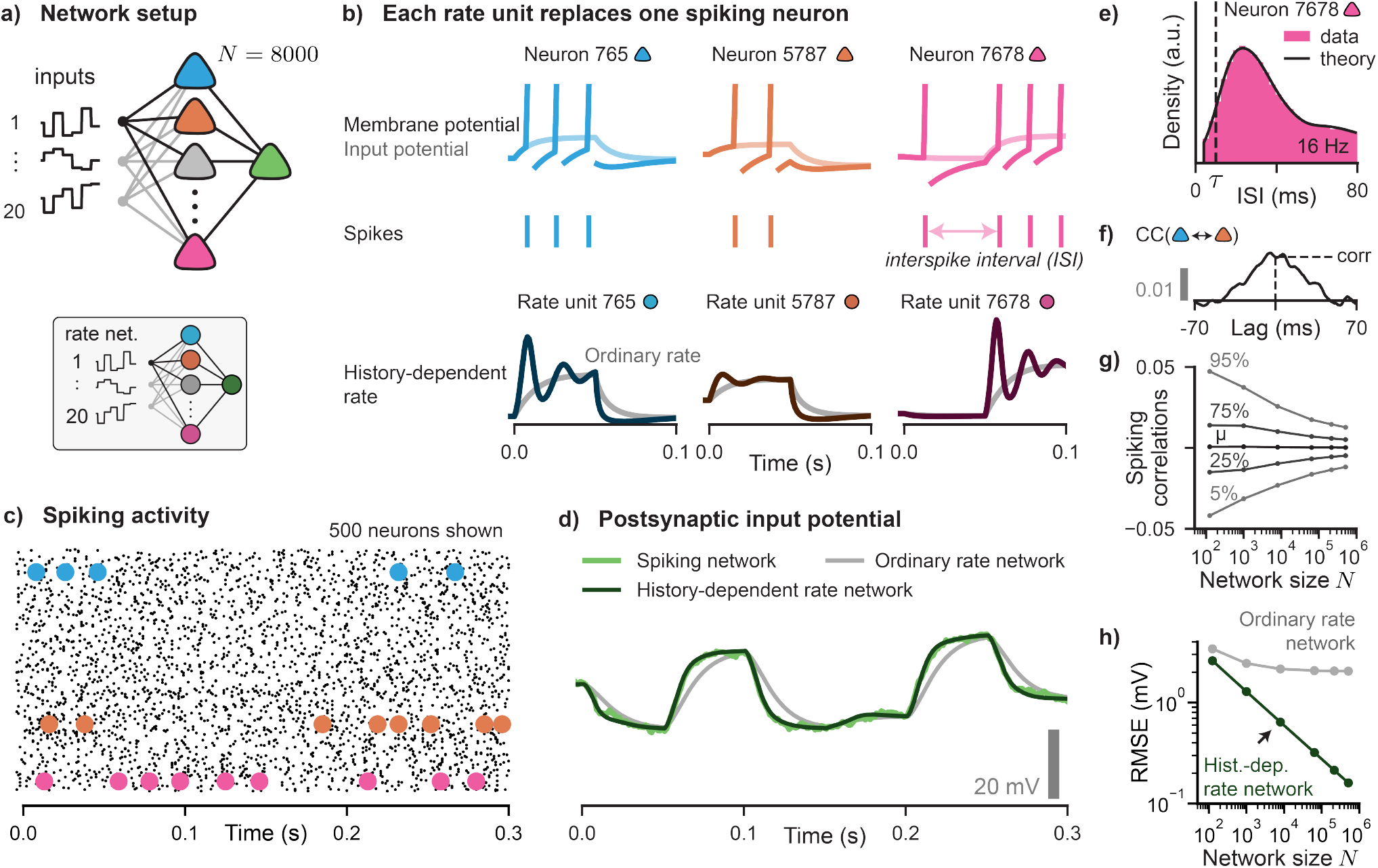
Heterogeneous spiking networks transmit fast temporal rate codes. **a)** Top: *N* = 8000 presynaptic SRM_0_ spiking neurons project to one postsynaptic neuron (green). The spiking neurons are driven by mixtures of *P* = 20 independent inputs. Bottom: A corresponding rate network with identical inputs and connectivity such that each rate unit replaces one spiking neuron. **b)** Dynamics and output of three example spiking neurons (top) and corresponding firing rate in the history-dependent rate network (bottom) with matching color code (blue, orange, pink). Vertical bars in the membrane potential (top) indicate spike emission, light colors show input potentials. History-dependent rate units (bottom) exhibit transient oscillations which are absent in the ordinary rate network (gray). Time measured from stimulus onset at *t* = −10 ms. **c)** Spike trains of 500 randomly selected neurons. Colored dots indicate spikes from neurons highlighted in b). **d)** Input potential of the postsynaptic neuron in the spiking network (green), history-dependent rate network (dark green) and ordinary rate network (gray). **e)** The interspike-interval (ISI) distribution of the pink spiking neuron in b); long simulation. Membrane time constant *τ* (vertical dashed line) shown for comparison. For theory, see Methods. **f)** Spike-spike cross-correlation (CC, Pearson) as a function of time lag between the blue and orange neuron in b), based on filtered spike trains. Dashed line indicates correlation at zero time lag. **g)** Distribution of spike-spike cross-correlations at zero lag between pairs of filtered neuronal spike trains (*µ*, mean; x%, x-th percentile; all pairs of *n* = 300 neurons randomly selected from the total of *N*) for varying network size *N* and number of external inputs *P* according to *P* = *N* ^1*/*3^. **h)** Root Mean Squared Error (RMSE) in the postsynaptic input potential (mean across *n* = 30 initializations) between spiking network and rate network, for both rate networks, for *P* = *N* ^1*/*3^. Arrow indicates network sizes in a-d.

Our mapping is highly accurate. The input potential *h*^out^(*t*) of the output neuron in the spiking network is predicted to a high degree of accuracy by the input potential *x*^out^(*t*) of the history-dependent rate network (Fig. 2d), as quantified by a small Root Mean Squared Error (RMSE) of less than 1 mV (Fig. 2h, arrow). For comparison, we also constructed a corresponding ordinary rate network and drove it with the same external input as the spiking network. In the ordinary rate network, the resulting output signal only approximates *h*^out^(*t*) well at time points where the external input has been constant for some time (Fig. 2d), leading to an RMSE which is more than 3 times larger than that of the history-dependent rate network (Fig. 2h, arrow; comparison of means across *n* = 30 initializations).

Looking at individual responses, we find that the history-dependent rate network predicts instantaneous firing rates that resemble the spiking patterns of the neurons they replace (Fig. 2b). For a spiking neuron *i*, a sudden increase in the input causes a rapid first spike to be emitted, followed by somewhat regular interspike intervals (ISIs) due to the refractoriness of the neuron (Fig. 2b, top). The same external input applied to rate unit *i* elicits a rapid peak in the firing rate, followed by a transient oscillation with a period comparable to the spiking intervals of neuron *i* (Fig. 2b, bottom). This refractory-like effect – absent in the ordinary rate network – can be directly attributed to the history-dependence of our rate equations and is shaped by the neuronal parameters and the received inputs (Gerstner, 2000).

Our mapping was successful even though network dynamics were fast and neurons are not Poisson-like. The frequent abrupt changes in the external input made spiking neuron responses more reliable than Poisson processes. To confirm this, we drove the spiking network with 50 repetitions of the same step-like external input (2000 ms each), and found that neurons had reliable spike counts per repetition (Fano factor of 0.19 ± 0.07; mean ± s.d. across *N* = 8000 neurons) and relatively regular ISIs (coefficient of variation of 0.79 ± 0.11), indicating highly non-Poisson behavior. Additionally, the ability to construct an accurate mapping could not be explained by strong low-pass filtering at the postsynaptic neuron, as neurons exhibited moderate firing rates (19 ± 8 Hz) and most ISIs were longer than the membrane time constant (96 ± 4 %; compared per neuron), causing characteristic ISI distributions with few intervals near zero due to refractoriness after spike emission (e.g. Fig. 2e; long simulation without repetitions). Instead, the accuracy of our mapping (Fig. 2d,h) is best understood through the concentration of measure phenomenon in large networks (Methods).

### Exact mapping for large one-layer networks

For one-layer networks as above, the mapping from a spiking network to a history-dependent rate network can be proven to be exact in the limit of large networks under suitable scaling of synaptic weights, even when no two neurons are alike (see Supplementary Note 1 in SI). To numerically verify this, we ran a series of network simulations of increasing network sizes, from *N* = 125 to *N* = 512, 000 presynaptic spiking neurons, where we simultaneously increased the number *P* of independent input signals driving the presynaptic neurons, according to *P* = *N* ^1*/*3^. This scaling ensures that spike-based input potentials concentrate to their mean, while simultaneously making networks more heterogeneous with an increase in the number of neurons (Schmutz et al., 2025). The growing heterogeneity can be understood by considering that, as network size grows, the total number 2^*P*^ of possible stimulus tunings that any neuron can have grows much faster than the total number *N* of neurons. Indeed, driving spiking networks with step-like inputs as before, we found that spiking correlations between any pair of neurons decrease as network size grows (Fig. 2g).

For each spiking network we constructed the corresponding history-dependent rate network. Looking at the input potential of the postsynaptic neuron, i.e., the network output which reads out one of the *P* signals from the presynaptic activity, we find that the RMSE between the spiking network output *h*^out^(*t*) and our history-dependent rate network output *x*^out^(*t*) decreases linearly as the number *N* of neurons in the network increases (Fig. 2h). In contrast, when comparing each spiking network to the corresponding ordinary rate networks, the RMSE between the two network outputs saturates at a level above 2 mV as network size grows, indicating a failure to accurately describe all aspects of neural communication in the spiking network. These results confirm that the effect of spiking variance on a postsynaptic neuron is reduced with the number of presynaptic input connections (despite increased heterogeneity), resulting in an effective time-varying drive to the postsynaptic neuron that can be deterministically described by the underlying instantaneous firing rates of the presynaptic neurons.

### History-dependent rate network for rapid signaling in multi-layer spiking networks

Multi-layer networks are particularly challenging to describe with rate-based networks since firing rates in one layer serve as inputs for the subsequent layer (through the function *G*_*i*_, Eq. (1)), which could lead to systematic mismatches in downstream layers (Brette, 2015). To study this scenario, we consider a spiking network with *N* = 3 × 8000 heterogeneous SRM_0_ neurons (three layers of *N*_*L*_ = 8000 neurons each; Fig. 3a), and compare it to a corresponding history-dependent rate network. Neurons in the first layer are driven by a mixture of *P* = 20 independent white Gaussian input signals, causing rapidly changing network dynamics. Subsequent layers are connected by low-rank weight matrices to allow simultaneous propagation of all *P* signals through the network with little interference (Hopfield, 1982, 1984; Mastrogiuseppe and Ostojic, 2018), yielding heterogeneous stimulus tuning on top of the heterogeneity in neuronal parameters. After three layers of processing, the signals are read out by *P* = 20 output neurons through weighted averages of the spiking activity of layer-three neurons. The fast external drive in layer one, combined with the structured weights between subsequent layers, results in network dynamics in all layers evolving at a speed comparable to the timescale of neural refractoriness, and yields highly diverse neural responses.

Our mapping retains a high degree of accuracy across all three layers. As before, we map the spiking network to a rate network by replacing each spiking neuron with a corresponding history-dependent rate unit, while retaining connection weights and single neuronal parameters (Fig. 3a). Driven by the same rapidly changing external input, we find that the history-dependent rate network predicts well the input potentials of the spiking network (Fig. 3b, left; RMSE *<* 1 mV for all layers for a simulation of *T* = 1s). This suggests that the history-dependent rate network provides an accurate description of the neuronal communication in the spiking network. Indeed, the history-dependent rates of individual units resemble the spike trains they replace, with localized peaks of activity reflecting reliable spike timing (Fig. 3b, right).

**Figure 3.**
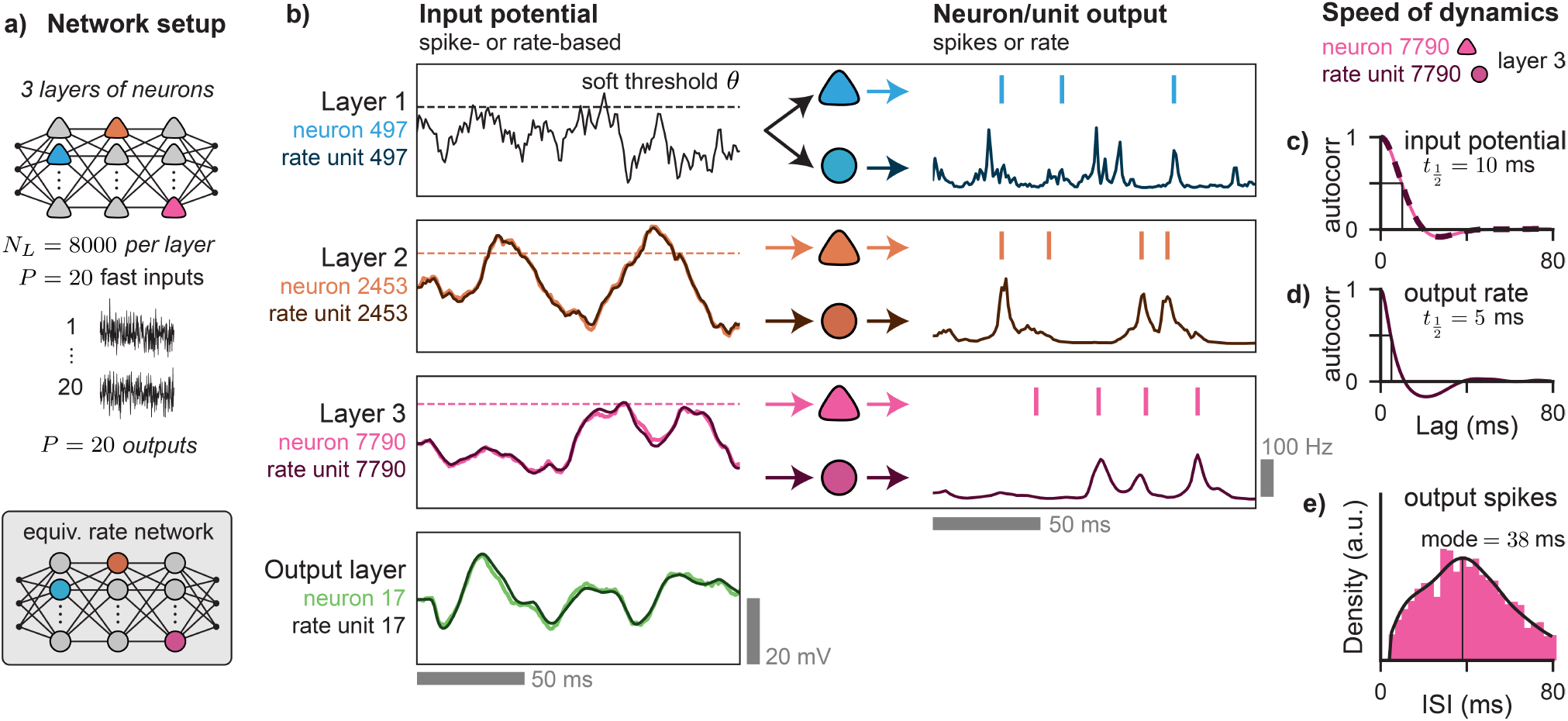
Fast rate coding across multiple layers. **a)** A feedforward network of SRM_0_ neurons driven by 20 white Gaussian signals is compared to the corresponding history-dependent rate network. **b)** Input-output relations for one example spiking neuron per layer (cyan, purple, pink) and corresponding rate units (darker colors). In the rate network, the rate output of one layer (right side) forms the input to the rate units of the next layer (left side, one row down). Network dynamics match those of the spiking network in all layers at the single-neuron level, including in the output layer (green). Layer 1 input potential (black) is identical for the spiking and rate network. **c-e)** Temporal statistics of the pink neuron in layer 3 and the corresponding rate unit (simulation of *T* = 100 s). **c**) Autocorrelation functions of the spike-based and rate-based input potential completely overlap. Black line indicates autocorrelation half life 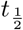. **d)** Autocorrelation function of the rate transmitted by the pink rate unit. **e)** Interspike-interval (ISI) distribution of the output spikes of the pink neuron. Black curve indicates theoretical estimate (Methods). Vertical line indicates predicted most likely interval.

Even in layer 3, the predicted rates are remarkably fast relative to the statistics of the spike trains they replace. We compared for each neuron-unit pair the autocorrelation of the predicted rate (e.g., Fig. 3d) to the interspike interval distribution of the spiking neuron it corresponds to (e.g., Fig. 3e), and found that predicted rates have mostly decorrelated by the time it typically takes for a neuron to fire a second spike. Across spiking neurons in layer 3, the most common interspike interval is 40.3 ± 10.2 ms long (mean ± s.d. across *N*_*L*_ = 8000 neurons; theoretical estimate per neuron, see Methods). However, the predicted rates of layer 3 rate units have a decorrelation half-time of 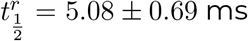 (mean ± s.d., *N*_*L*_ = 8000 units), which is shorter than 99.77 ± 0.64 % of corresponding ISIs in the spiking network (mean s.d. across *N*_*L*_ = 8000 neuron-unit pairs). This suggests that in the spiking network the underlying instantaneous firing rate of a spiking neuron changes substantially before a second spike is transmitted.

Our mapping is asymptotically exact for large multi-layer networks, and extends to networks of LIF neurons. We ran a series of simulations for both SRM_0_ and LIF networks for increasing sizes to examine the effect of network size on the accuracy of our mapping. For each spiking network we constructed the corresponding history-dependent rate network (Methods) and compared the neuron-level dynamics. We find that for both SRM_0_ and LIF networks, as the number *N*_*L*_ of neurons per layer increases, the mismatch between the spiking network and the corresponding history-dependent rate network decreases at the same constant rate in all layers (Fig. 4b,d), despite increasing heterogeneity due to the simultaneous increase in the number *P* of signals according to 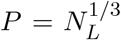. This shows that in large networks, for both SRM_0_ and LIF, neuronal communication becomes rate-like and near-deterministic despite the variable spike emission of individual neurons.

**Figure 4.**
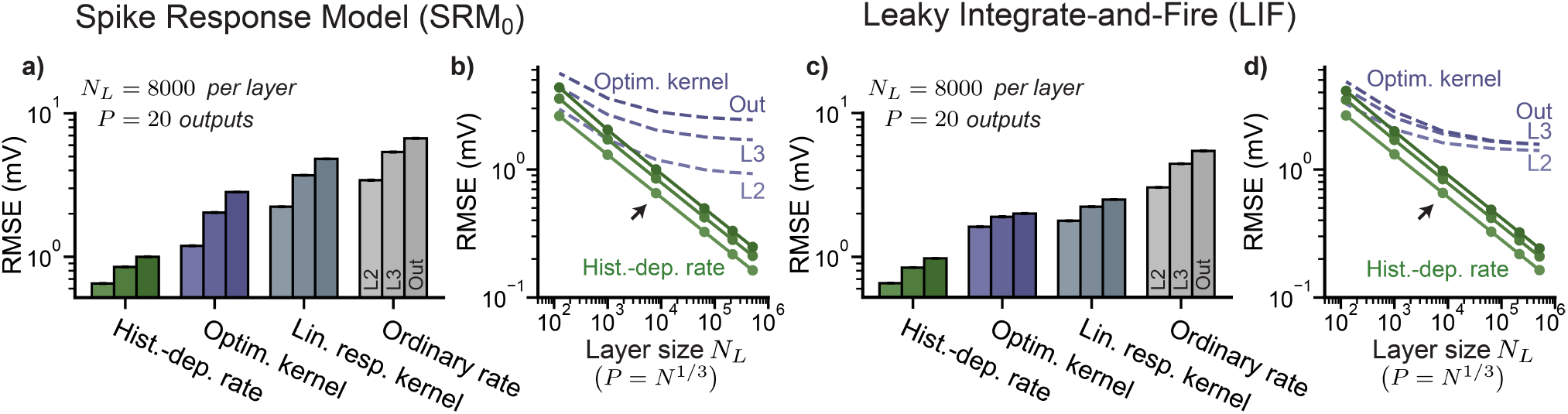
Vanishing mismatch as networks become large for both SRM_0_ and LIF. **a)** Mismatch compared to spiking network across layers (L2, L3, Out; lighter to darker colors) measured in RMSE (mean ± s.d. across *n* = 20 initializations) on a logarithmic scale, for corresponding rate networks constructed using our mapping (History-dependent rate) as well as three existing linear-nonlinear approaches (Optimized kernel; Linear response kernel; Ordinary rate). **b)** Mean RMSE (*n* = 15 initializations) across layers for varying number of neurons per layer *N*_*L*_ on a log-log scale, for the two best performing mappings (green, History-dependent rate; purple, Optimized kernel), using *P* = *N* ^1*/*3^. Arrow indicates network size in a). **c-d**) Same as a-b) for spiking networks consisting of Leaky Integrate-and-Fire neurons, compared to corresponding rate networks.

### Linear-nonlinear mappings cannot fully capture rich network dynamics

In our rate network, the explicit history dependence captures the rich short-term dynamics regardless of the speed or amplitude of the input potentials. To evaluate the importance of the history-dependence, we compared how well several existing linear-nonlinear rate equations can describe the fast dynamics of the SRM_0_ spiking network of the previous section and its LIF equivalent. In the linear-nonlinear rate networks, the firing rate equations take the form of a linear-nonlinear cascade (Hunter and Korenberg, 1986; Chichilnisky, 2001; Ostojic and Brunel, 2011), with the firing rate of rate unit *i* given by *r*_*i*_(*t*) = *F*_*i*_ ((*κ*_*i*_ * *I*_*i*_) (*t*)), where the rate-based input 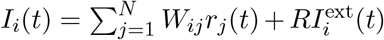 is first passed through a normalized, linear filter (kernel) *κ*_*i*_(*s*) that may be neuron-specific and then through the f-I curve *F*_*i*_(*x*) of spiking neuron *i* (Methods). Rate equations of this form ensure the correct stationary state and can capture some oscillatory behavior (Ostojic and Brunel, 2011).

We compared three kernel definitions (Methods), each leading to a different linear-nonlinear rate network. First, the optimized kernel approach consists of directly fitting the shape of a single kernel to be used for all units, optimized to reproduce the history-dependent rates of all units as accurately as possible for a select training stimulus. A method to fit per-unit kernels was also considered and yielded near-identical results to those of the shared kernel approach (Suppl. Fig. S1). Second, in the linear response kernel approach, each unit has a unique kernel inferred from the corresponding neuron’s response to a small pulse input (Ostojic and Brunel, 2011; Herrmann and Gerstner, 2001), with the linearization point optimized to reproduce the spiking network dynamics as accurately as possible. Finally, the ordinary rate equation is retrieved when an exponential kernel is used with decay rate set to the membrane time constant.

For each kernel definition, a corresponding rate network was constructed with identical architecture to that of the multi-layer SRM_0_ spiking network (*N* = 3 × 8000, *P* = 20), each driven with the same external input to the first layer as the spiking network. While both the optimized shared kernel approach and the linear response kernel approach lead to better neuron-level descriptions of the spiking network dynamics than the ordinary rate network, all linear-nonlinear rate networks suffer from considerable error accumulation across layers (Fig. 4a), making the final network output considerably different from that of the spiking network (Suppl. Fig. S2).

Even the best-performing linear-nonlinear mapping, the optimized shared kernel approach, is not asymptotically optimal. When using this mapping to construct rate networks for spiking networks of varying sizes, we find that the mismatch between outputs of the spiking network and the corresponding linear-nonlinear rate network saturates at a value above 2 mV as the number *N*_*L*_ of neurons per layer increases (Fig. 4b). Similar results were obtained for LIF spiking networks (Fig. 4c-d). Together, these results suggest that no linear-nonlinear rate network can fully describe the neuronal communication in renewal-type spiking networks, and highlight the importance of an accurate rate model when describing networks with multiple layers of processing.

### Reduced correlations in sparse networks improve rate coding

Spiking variability in presynaptic neurons produces small, unpredictable fluctuations in postsynaptic input potentials. For the fully connected feedforward architecture considered so far, the amplitude of spiking-induced fluctuations is found to scale proportional to 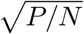 for Poisson neurons (Schmutz et al., 2025). However, biological networks are not fully connected but show sparse connectivity. We investigate how sparsity, quantified by the number of incoming connections *K < N* per neuron, influences our mapping.

We studied two reference spiking networks each with three layers of SRM_0_ neurons, one with *N* = 3 × 8000, *P* = 20 and one with *N* = 3 × 16000, *P* = 10. For each reference network we systematically decreased the number of incoming connections per neuron from fully connected (*K* = *N*_*L*_) to sparse 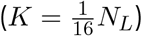 while rescaling synaptic strengths accordingly. When neuron-level dynamics are compared with corresponding history-dependent rate networks, we find that the RMSE in layer 2 is comparable to that found in fully connected networks with size *N*_*L*_ = *K* (Fig. 5a), and scales proportional to the fraction 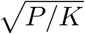. In deeper layers, the RMSE also scales proportional 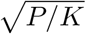, but for any specific network has a smaller value than for a fully connected network with the same number of incoming connections per neuron.

**Figure 5.**
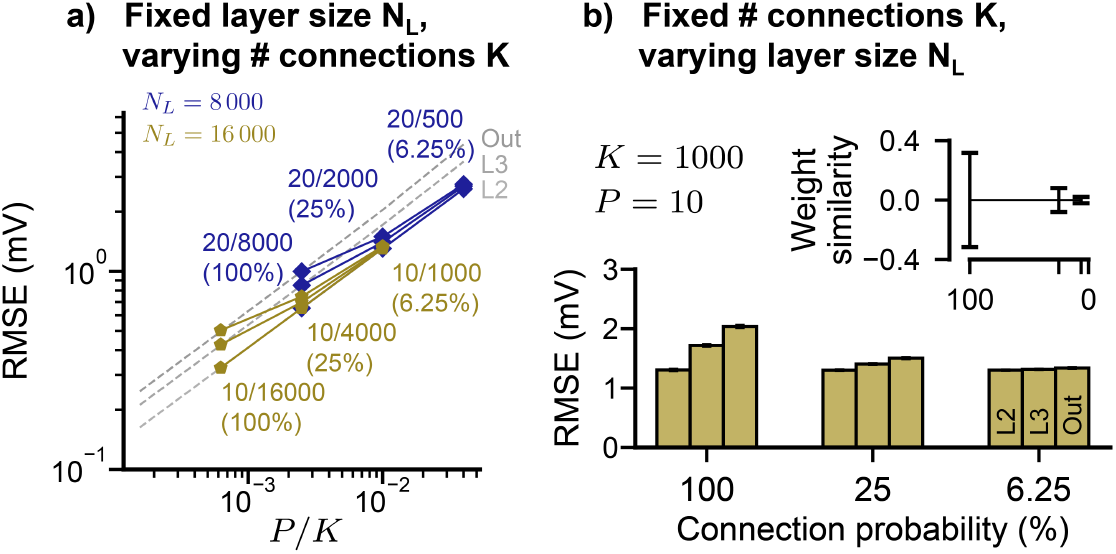
Sparsity reduces mismatch accumulation. **a)** Mismatch between the history-dependent rate network and spiking network (RMSE across *n* = 150 network initializations; for layers 2, 3 and Out) as a function of *P/K*, where *K* is the number of incoming connections per neuron, for fixed number *N*_*L*_ of neurons per layer (blue, 8000; yellow, 16 000). Gray, dashed lines repeat fully-connected network in Fig. 4b. **b)** For fixed number *K* of incoming connections, mismatch accumulation from layer-to-layer decreases with lower connection probability *K/N*_*L*_. Inset: distribution of pairwise cosine similarity between vectors of incoming connection weights as a function of connection probability (errorbars, mean ± s.d.; pairs from *n* = 1000 neurons randomly selected from the total).

To further investigate this observation, we ran another series of simulations in which we increased the layer size *N*_*L*_ while keeping the number of incoming connections as well as the number of inputs fixed (*K* = 1000, *P* = 10), ensuring *P/K* remains fixed while sparsity is varied from fully connected (*N*_*L*_ = *K*) to sparse (*N*_*L*_ = 16*K*). First, we find that the mismatch in layer 2 is not affected by the increasing layer size, consistent with the scaling law found above. Second, increasing sparsity while keeping *K* fixed decreases the RMSE in deeper layers (Fig. 5b), with the RMSE in all deeper layers approaching that of layer 2. This can be understood as the result of reduced similarity in presynaptic connections between any pair of neurons (Fig. 5b, inset), which causes decreased spiking correlations and less accumulation of spiking fluctuations across layers (Brunel, 2000; Tetzlaff et al., 2003; Vogels and Abbott, 2005; Kumar et al., 2010; Brette, 2015).

## Discussion

The conditions under which mappings from spikes to rates are possible also provide conditions under which rate coding is meaningful in biological data (Brette, 2015). This work shows that the dynamics of noisy spiking networks can be accurately reduced to rate-based interactions, even if similarly tuned neurons are absent, dynamics are fast, and neuronal firing is non-Poisson. Under the conditions on the connectivity identified in this paper, a postsynaptic neuron “sees” the instantaneous firing rates of its presynaptic neurons. Commonly used rate models such as ordinary rate networks or more general linear-nonlinear rate networks are unable to capture the rich dynamics of fast rate codes, whereas our historydependent rate network yields an accurate description of neuronal communication in large feedforward networks able to process many signals in parallel.

Fast rate coding has long been known for homogeneous populations of Integrate-and-Fire neurons (Knight, 1972; Gerstner, 2000; Brunel et al., 2001), as they can encode signals that vary faster than the neurons’ membrane time constant. Our theory generalizes fast rate coding to heterogeneous network where each neuron has different parameters and a different stimulus tuning. Somewhat unexpectedly, the mathematical formulation of a single history-dependent rate unit is identical to that of a homogeneous population, which means that the mathematical tools known for homogeneous populations (Gerstner, 2000; Brunel et al., 2001; Fourcaud and Brunel, 2002) can be applied to networks with a large degree of neuronal heterogeneity (Gast et al., 2024). Consistent with the theories of fast rate coding, neuronal recordings *in vitro* and *in vivo* have shown that single neurons can reliably encode time-varying input signals in their peri-stimulus time histogram (PSTH) up to frequencies of ~200 Hz (Köndgen et al., 2008; Tchumatchenko et al., 2011), which is much faster than what passive membrane potential dynamics would allow.

Our theory of fast rate coding has at least three implications for the analysis of neural data. Firstly, exploiting the link between rate coding and latent population-level factors (DePasquale et al., 2023), the theory of fast rate coding should encourage neuroscientists to search for fast latent factors, for example, by reducing the time scale parameter (Yu et al., 2009; Macke et al., 2011; Aghagolzadeh and Truccolo, 2014; Pandarinath et al., 2018) or by including history-dependence (Wang et al., 2022). Secondly, the existence of an exact rate formalism with smooth rate variables opens the door to fitting more realistic spiking networks to largescale data (Urai et al., 2022) using smooth gradient methods for measured as well as ‘hidden’ neurons without sampling spikes (Bellec et al., 2021; Sourmpis et al., 2026). Thirdly, the existence of an exact expression for the time-dependent firing rate should facilitate parameter fitting under the constraint of correct trial-averaged activity (Bellec et al., 2021; Sourmpis et al., 2026).

Our theory of fast rate coding also has implications for the optimization of (recurrent) spiking neural networks. While the dynamics of RNNs with ad hoc rate equations may transfer to spiking networks (Kim et al., 2019; DePasquale et al., 2023), both the naturally slow dynamics of RNNs as well as the simplified rate description may limit which solutions can be found. Instead, the exact equivalence between spikes and smooth firing rates could allow for gradient-based optimization on history-dependent rate networks, the solution of which could then be transferred to an equivalent spiking network. The ability to derive exact gradients may also pose as an alternative to surrogate gradients (Neftci et al., 2019), which are already related to noisy spike generation (Gygax and Zenke, 2025). Similarly, the generalized rate formalism may allow FORCE-based approaches to be extended beyond the quasistationary approximation, allowing them to capture fast target dynamics by defining errors using accurate estimates of instantaneous firing rates given spike-based inputs (Kim and Chow, 2018) instead of low-passed spikes, as is more common (Abbott et al., 2016; Nicola and Clopath, 2017; DePasquale et al., 2023; Newton and Nicola, 2025).

The parallel transmission of *P* signals in a network of *N* ≫ *P* neurons is made possible by low-rank feedforward weight matrices. Low-rank connectivity is commonly used in computational neuroscience (Mastrogiuseppe and Ostojic, 2018; Pezon et al., 2026). Moreover, task-trained RNNs often have weight matrices dominated by a low-rank component (Sussillo and Barak, 2013; Dubreuil et al., 2022; DePasquale et al., 2023). In our theory, the rank *P* is relatively large (up to 80 in simulations) and the RMSE between spiking and the rate networks scales as 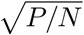 (see Supplementary Note 1 in SI). While we have considered the scaling *P* = *N* ^1*/*3^ for our simulations, our theory predicts that even in the case where synapses are strong, so that each weight vector has a length of order one (Rubin et al., 2017) and *P* grows linearly with *N*, the RMSE between spiking and rate networks should remain small as long as the ratio *P/N* is small.

We believe that the general principles underlying fast rate coding through virtual populations should extend to networks of any type of spiking model, such as spiking neuron models with input noise (Brunel and Hakim, 1999; Brunel, 2000), adaptation (Muller et al., 2007; Schmutz, 2022), non-linear membranes (Fourcaud-Trocmé et al., 2003), or combinations thereof (Brette and Gerstner, 2005). While for those advanced spiking models the simulation of the corresponding rate models would be costly than our history-dependent rate network, it is important to know that rate representations of these models do exist. Further, if we tolerate some mismatch, approximations of the exactly equivalent rate model can be used, such as linear-nonlinear cascades (Ostojic and Brunel, 2011), eigenmode decomposition (Pietras et al., 2020), or quasi-renewal theory (Naud and Gerstner, 2012). The latter includes both short-term effects caused by refractoriness and long-term effects caused by adaptation, whereas earlier history-dependent rate equations only cover adaptation (Benda and Herz, 2003). Additionally, while we have focused on noise-free input potentials in our rate network, the inclusion of finite-size corrections to population equations (Schwalger et al., 2017; Mattia and Del Giudice, 2002; Vinci et al., 2023) may help improve rate descriptions in smaller networks with heterogeneity.

The instantaneous firing rate of one rate unit in our network could potentially also be interpreted as the trial-averaged spike trains of a single spiking neuron in experiments under the assumptions of our mapping. One important assumption is that this requires any observed trial-to-trial variability to be purely noise which does not affect downstream neural processing, thereby excluding cases of shared spiking variability caused by stimulus-independent ongoing activity (Arieli et al., 1996; Stringer et al., 2019). When this holds, our work implies that the trial-averaged activity of a single neuron could represent the signal that this neuron communicates to other neurons in any single trial, despite the fact that the trial-averaged activity is not observable in a single trial. This generalizes the classic PSTH rate coding argument made for pools of similarly tuned neurons (Perkel and Bullock, 1968; Shadlen and Newsome, 1994). From a more general perspective, our model of rate coding in heterogeneous networks offers a theoretical foundation for the widespread use of rate models in computational neuroscience as well as for rate-based analysis of experimental data.

## Acknowledgments

This work was supported by the Swiss National Science Foundation grants 200020-207426 and 200021-236436. V.S. is supported by a fellowship from the Swiss National Science Foundation (grant no. 222150).

## Methods

### The ordinary rate network and linear-nonlinear generalizations

A standard recurrent neural network is a special case of rate networks as defined in Eq. (1). To see this, let us assume that functions *G*_*i*_ take the specific form of a linear-nonlinear cascade (Hunter and Korenberg, 1986; Chichilnisky, 2001; Simoncelli et al., 2004; Herrmann and Gerstner, 2001; Ostojic and Brunel, 2011). Then, the rate network takes the familiar form,

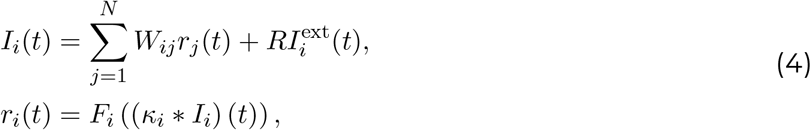

where *κ*_*i*_ is a normalized causal kernel, i.e., ⎰*κ*(*s*)*ds* = 1 and *κ*(*s*) = 0 for *s* 0. The total input *I*_*i*_(*t*) unit *i* receives, which is the sum of synaptic inputs and the external input, first passes through the linear filter *κ*_*i*_ and then through an instantaneous (i.e., time-local) nonlinearity *F*_*i*_. If all the *κ*_*i*_’s are exponential filters, 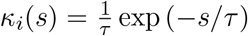, the equations above reduce to the well-known differential equations of an RNN,

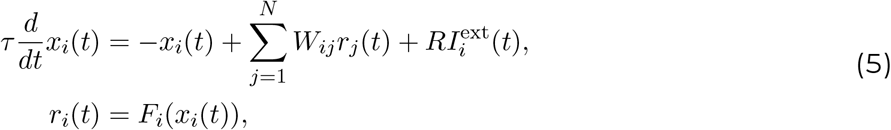

as explained, e.g., in (Dayan and Abbott, 2001, Chapter 7.2) and (Gerstner and Kistler, 2002, Chapter 6.3).

The *ordinary rate network* is obtained by mapping a spiking network one-to-one onto a rate network in the form of Eq. (5), keeping the same connectivity *W*_*ij*_ and setting each *F*_*i*_ as the ‘f-I curve’ of spiking neuron *i*, i.e., the function that gives the steady-state firing frequency of a neuron as a function of the constant input it receives (see Eqs. (25) and (26)).

The more general *linear-nonlinear rate networks* we consider keep the nonlinearity *F*_*i*_ fixed to the f-I curve of spiking neuron *i*, but allow for arbitrary, possibly neuron-specific, linear filters *κ*_*i*_. The use of the f-I curve ensures the correct stationary state, while the kernel shape can capture transient components of the response (Ostojic and Brunel, 2011). While models of this type are numerically and analytically convenient, the mapping from a spiking network to a linear-nonlinear rate network is only accurate when spiking neurons are linear-nonlinear Poisson neurons or when network dynamics are slow enough to be considered quasi-stationary.

### Spike response model with heterogeneous parameters

An SRM_0_ neuron takes into account refractory effects following spike emission. After the emission of a spike from neuron *i*, the neuron has to recover before it can emit another spike. Both the absolute and relative refractoriness are modeled through the (negative) *refractory kernel* 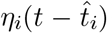, where 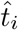 is the most recent firing time of the neuron *i*. The *membrane potential u*_*i*_(*t*) of a neuron *i* is then the combination of the input potential *h*_*i*_(*t*) of Eq. (2) of the main text with the refractory kernel, i.e., 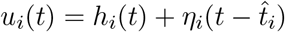. The membrane potential *u*_*i*_(*t*) determines the probability for neuron *i* to emit a spike in a short time window around time *t* through a neuron-specific soft-threshold function *f*_*i*_ which increases with the membrane potential and is typically negligible below a certain threshold. At any given time *t*, the probability for neuron *i* to emit a spike in the small time interval from *t* to *t* + *Δt*, conditioned on the input potential *h*_*i*_(*t*) and and the last firing time 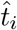, is given by

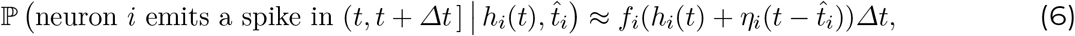

where the approximation tends to an equality as *Δt* tends to 0. Formally, spiking times are generated by a time-dependent renewal process with instantaneous firing intensity given by 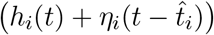, and spike current pulses are modeled with the Dirac delta function. Together, Eqs. (2) and (6) define the dynamics of the spiking neural network. Throughout this paper we assume that the spike current pulse is a Dirac delta function, i.e., integration 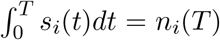 yields the number of spikes in the interval from 0 to *T*. In other words *n*_*i*_(*T*) is a counting process as a function of *T*.

For the functions *f*_*i*_ and *η*_*i*_ we choose

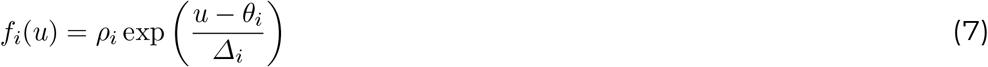

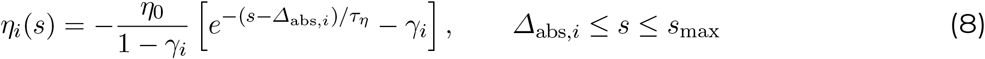

with a finite history dependence, *η*_*i*_(*s*) = 0 for *s* ≥ *s*_max_, and with an absolute refractory period modeled as *η*_*i*_(*s*) = −∞, for 0 *< s < Δ*_abs,*i*_. The constant 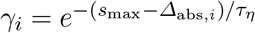 offsets and normalizes the kernel such that *η*_*i*_(*Δ*_abs,*i*_) = −*η*_0_ and *η*_*i*_(*s*_max_) = 0. The argument of the exponential in *f*_*i*_(*u*) is clipped to be at most 5 to ensure numerical stability.

In the simulations, we allow for a neuron-specific resting potential *u*_rest,*i*_; however, in the equations above, the voltage *u* and the neuron-specific threshold *θ*_*i*_ are measured relative to rest to simplify the notation. This is possible, because of the shift-invariance of Eq. (7), i.e., only the difference *u* − *θ*_*i*_ matters. At initialization, for each neuron *i* the last spiking time 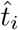 is sampled from the neuron’s theoretical stationary ISI distribution. Heterogeneous neuron types are obtained by sampling neuron parameters independently from probability distributions listed in Tab. 1.

**Table 1.**
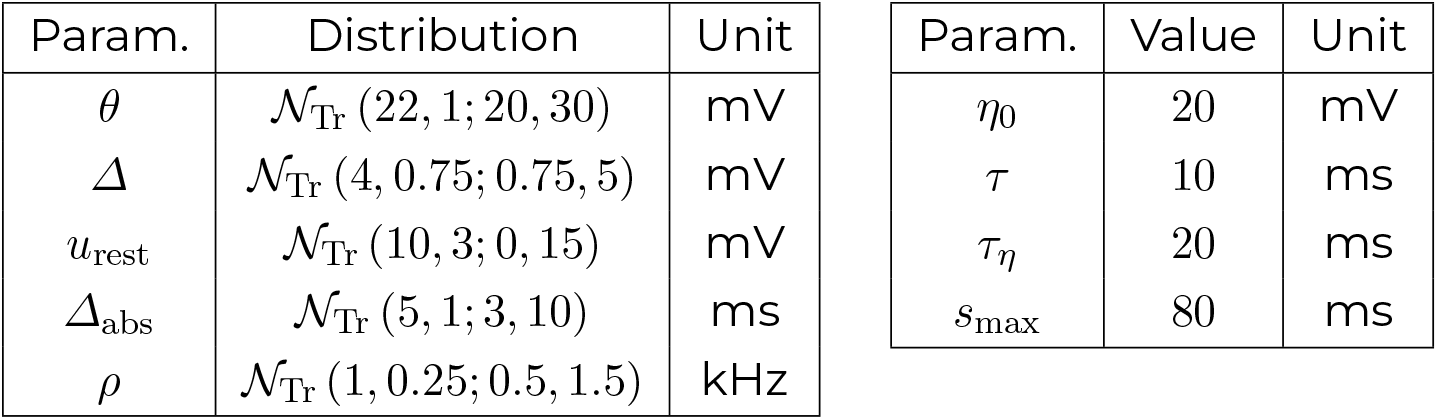
Heterogeneous SRM_0_ parameters used in simulations unless mentioned otherwise. *N*_Tr_ (*M, σ*; *a, b*) is the truncated normal distribution with mean *M*, standard deviation *σ*, and support [*a, b*]. Note that the threshold *θ* indicated in the table represents the absolute threshold used in simulation. Notation in the text was simplified by exploiting the shift-invariance between *u*_rest_ and *θ*, meaning each *θ*_*i*_ and *u*_*i*_(*t*) in the text should be seen as measured relative to the resting potential.

| Param. | Distribution | Unit | Param. | Value | Unit |
|---|---|---|---|---|---|
| θ | 𝒩<sub>Tr</sub> (22, 1; 20, 30) | mV | η<sub>0</sub> | 20 | mV |
| Δ | 𝒩<sub>Tr</sub> (4, 0.75; 0.75, 5) | mV | τ | 10 | ms |
| u<sub>rest</sub> | 𝒩<sub>Tr</sub> (10, 3; 0, 15) | mV | τ<sub>η</sub> | 20 | ms |
| Δ<sub>abs</sub> | 𝒩<sub>Tr</sub> (5, 1; 3, 10) | ms | s<sub>max</sub> | 80 | ms |
| ρ | 𝒩<sub>Tr</sub> (1, 0.25; 0.5, 1.5) | kHz |  |  |  |

### Spiking statistics

Long simulations (*T* = 2 hours) were run for select neurons to determine statistics of interest, such as the inter-spike interval distribution and pairwise spiking correlations. Spiking cross-correlations (CCs) were computed at various time lags as the Pearson correlation between two filtered spike trains, each convolved with a rectangular filter of width 5 ms (Renart et al., 2010). Equivalent simulations were run for corresponding history-dependent rate units to find theoretical ISI distributions.

### Measure of neuron-level mapping accuracy

Given a spiking network and a rate network driven by the same external input, the neuron-level mismatch in the dynamics is calculated using the Root Mean Squared Error (RMSE). First, the MSE (without the square root) is calculated between the input potential of neuron *i* in layer *l* in the spiking network, 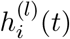, and the corresponding input potential of rate unit *I* in the rate network, 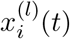,

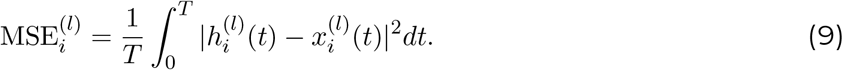

for a simulation time *T* = 10 000 ms unless indicated otherwise.

The RMSE of a layer (*l*) is the square root of the average MSE in that layer,

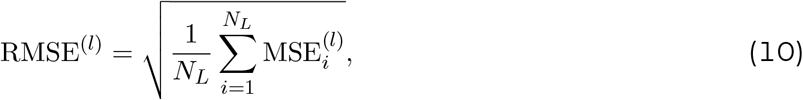

where the average is taken over all neurons/units in that layer (*N*_*L*_ for intermediate layers, *P* for output layer).

### History-dependent rate network through virtual populations

To derive rate networks that generalize linear-nonlinear rate networks, we imagine that we replace each spiking neuron by a virtual population of identical neurons. The populations are virtual in the sense that they serve only as a device for deriving a rate network; the mapping from spiking to rate is neuron-to-unit (not population-to-unit), and can be applied to heterogeneous spiking networks (without duplicate neurons).

The virtual population approach consists of imagining that each neuron *i* in the spiking network is replaced by a virtual, infinite population of identical copies of neuron *i*; sharing the same neuronal parameters of neuron *i* and all receiving the same input, but with individual variability emerging from stochastic spike emission through escape noise (Eq. 6). For each rate unit *i* in the rate network, the firing rate *r*_*i*_(*t*) is defined by the input-output relation of this virtual population. That is, the function *G*_*i*_ (Eq. 1) is defined by the population-averaged spiking activity in response to a (rate-based) input potential *x*_*i*_(*t*).

For a network of SRM_0_ neurons, the virtual population approach leads to a rate network where the firing rate *r*_*i*_(*t*) of each rate unit *i* is described by an integral equation (Gerstner, 2000).

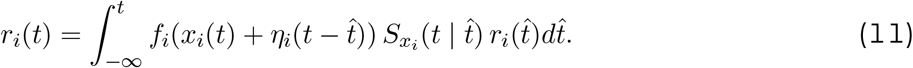

which, together with the equation for the rate-based input potential *x*_*i*_(*t*) (Eq. 3), defines the corresponding history-dependent rate network. This model is “history-dependent” in the sense that the rate *r*_*i*_(*t*) depends on its own history, and it takes into account neuronal refractoriness by retaining the refractory kernel *η*_*i*_ of the corresponding spiking neuron, allowing it to capture fast transient responses.

The function 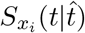 in Eq. (11) is the survivor function of renewal theory (Cox, 1962) extended to time-dependent input (Gerstner, 2000), the subscript *x*_*i*_ being a reminder that the probability, 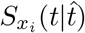, to have “survived” without firing from time 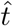 to *t* depends on the rate-based input potential *x*_*i*_(*t*^*′*^) on the time interval 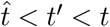.

For networks of LIF neurons, the same virtual population approach leads to a similar history-dependent rate network (see Eq. 17).

### Formal derivation of the history-dependent rate network

To formally arrive at Eq. 11, we use that the population-averaged firing rate of a virtual population is identical to the expected instantaneous firing rate of a single neuron, conditioned on its past input potential. The number of spikes emitted by a neuron *i* over the time interval (*t, t* + *Δt*] is given by the integral over the spike train

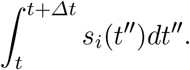

Hence, the expected instantaneous firing rate of neuron *i* conditioned on its past input potential is 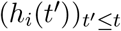 is

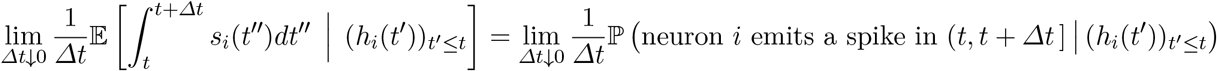

If neuron *i* is an SRM_0_ neuron, the conditional expected instantaneous firing rate is given by

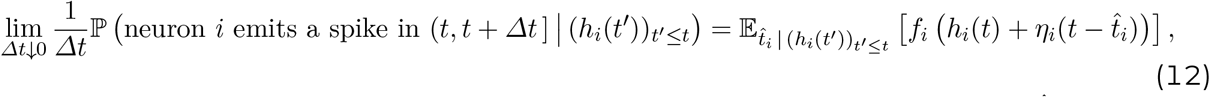

where, on the right-hand side, the expectation is taken over the last spike time 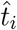 of neuron *I* conditioned on the past input potential 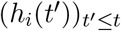.

To obtain a rate network from the expression on the right-hand side of Eq. (12), we replace the spike-based input potential *h*_*i*_(*t*) in Eq. (12) by its rate-based counterpart *x*_*i*_(*t*) (see Eq. (3)); this yields the purely rate-based expression for the instantaneous firing rate of neuron *i*,

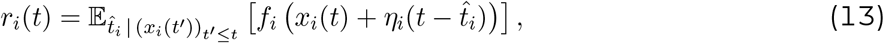

where the expectation is taken over the time of last spike 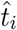 of an SRM_0_ neuron, whose past input potential is 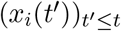 instead of the original 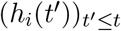. By using *r*_*i*_(*t*) defined by Eq. (13) in the equation for the rate-based input potential *x*_*i*_(*t*) in Eq. 3, the system of equations is fully closed in terms of firing rates, thereby defining a general rate network in the sense of Eq. (1) since the firing rate *r*_*i*_(*t*) only depends on the past firing rates of the other rate units (through the rate-based input potential *x*_*i*_(*t*)).

Time-dependent renewal theory tells us how the expectation on the right-hand side of Eq. (13) can be expressed as an integral over unit *i*’s past rate (Gerstner, 2000). For renewal-type neurons, the probability distribution of the last spike time 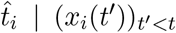 can be written as 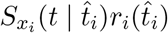, where 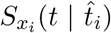 is the survivor function. In the case of a SRM_0_ neuron, the survivor function is

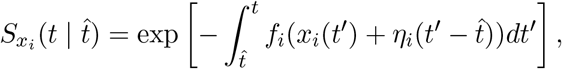

which completes the derivation of the integral equation for the rate *r*_*i*_(*t*), Eq. (11).

### Networks of Leaky Integrate-and-Fire neurons

An LIF neuron *i* is typically formulated directly through a membrane potential *u*_*i*_(*t*), that resets to a fixed value upon spike emission:

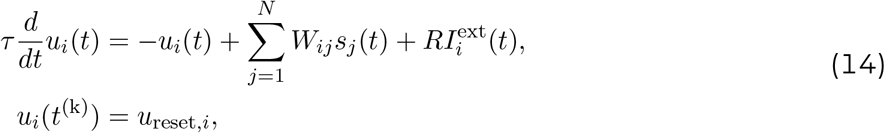

following each spiking time *t* = *t*^(*k*)^, with dynamics of *u*_*i*_(*t*) disabled during an absolute refractory period of duration *Δ*_abs,*i*_ following spike emission. Spikes are generated stochastically with escape noise, with an instantaneous firing intensity *f*_*i*_(*u*_*i*_(*t*)) as in Eq. (7).

The LIF model above can be formulated as a variant of the SRM_0_, by describing the neuron by an input potential *h*_*i*_(*t*) (Eq. 2), and writing the membrane potential *u*_*i*_(*t*) as a sum of *h*_*i*_(*t*) and a refractory kernel that now depends on the input potential at the time of reset,

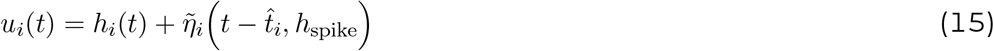

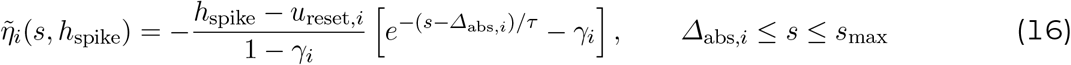

where 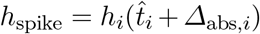 is the *input* potential at the time when the neuron starts integrating inputs again, right after the absolute refractory period *Δ*_abs,*i*_ ends. The refractory time constant is equal to the membrane time constant *τ*. If the absolute refractory period is short and the noise level is low, the typical value of *h*_spike_ is *h*_spike_ ≈ *θ*_*i*_. Additionally, *u*_reset,*i*_ determines the *membrane* potential after reset. This formulation is identical to Eq. (14) up to the finite size of the refractory kernel, and becomes exact for *s*_max_ → ∞.

To create a heterogeneous spiking network with LIF neurons, neuron parameters in simulations are randomly sampled from the same neuron parameter distributions as for SRM_0_ (Tab. 1) except for *u*_rest,*i*_ ~ N _Tr_ (7, 3; 0, 15). Additionally, *u*_reset,*i*_ ~ N _Tr_ (−2, 3; −10, 0). Distributions are chosen to produce similar mean firing rates as SRM_0_ neurons.

From this the corresponding history-dependent network can be derived in a similar selfconsistent form as before.

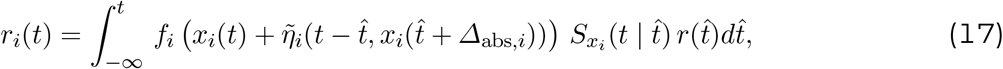

where 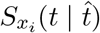 is the survivor function,

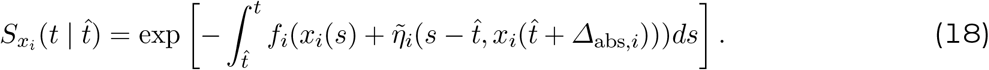

### Network connectivity for structured information processing

Structured connectivity is used to impose meaningful information processing. Feedforward networks are constructed to robustly process a small number *P* of signals (*P < N*_*L*_) simultaneously, with minimal interference between signals. This is implemented through a low-rank framework with layer-to-layer connectivity matrices following a Hopfield-like structure (Hopfield, 1982). Each of the *P* signals corresponds to a pattern index *µ*, and weights are constructed by a combination of binary variables 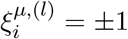 (i.i.d. sampled Rademacher variables) where *l* is the layer index. Specifically, the connection weight from neuron *j* in layer *l* − 1 to neuron *i* in layer *l* is

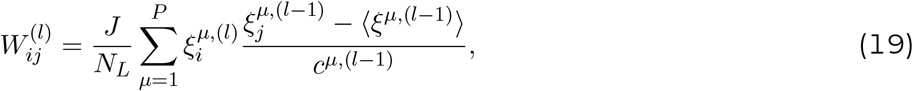

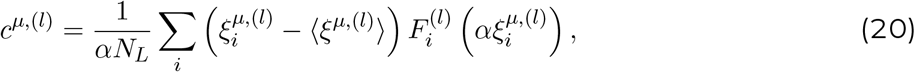

where 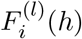 is the neuron-specific stationary firing rate (i.e., f-I curve) at a stationary input potential *h*, and *α* determines the desired scale of the input potentials in response to the signals. Brackets denote an average over neurons in a layer, i.e., 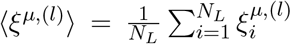. To ensure approximately constant signal strength from layer to layer, weights are divided by the normalization constants *c*^*µ*,(*l*)^ and multiplied with an empirical factor *J* of order one for fine-tuning, ranging from *J* = 0.71 to *J* = 1.4 depending on the type of external input (step-like or white noise) and the neuron model used (SRM_0_ or LIF).

External input consists of *P* independent input currents *I*_*µ*_(*t*) that drive neurons in the first layer along the corresponding patterns, i.e., 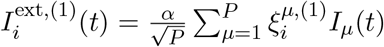. Neurons in deeper layers receive no external input. Network output consists of *P* linear readout neurons, each extracting the respective signal from neurons in layer *L*.

We considered two types of external input. First, *P* independent step-like inputs that change every 50 ms, with for each pattern *µ*, a new step height sampled uniformly from −1 to 1 every interval. For this input type we used *α* = 14 mV and *J* = 1. Second, *P* independent white Gaussian inputs, such that the input potential of any neuron in the first layer becomes an independent mixture of *P* Ornstein-Uhlenbeck processes. Here we used *α* = 30 mV and *J* = 1.4 or *J* = 0.71 for SRM_0_ and LIF, respectively.

### Efficient discrete-time implementation of the history-dependent rate network

Numerical forward integration of the spiking network dynamics (Eq. 2) is performed using the forward Euler method

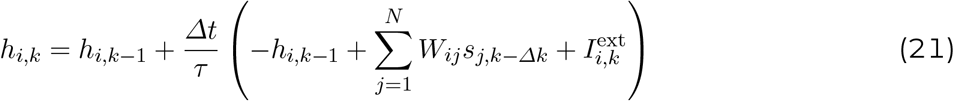

where *h*_*i,k*_ = *h*_*i*_(*kΔt*) with discretization time *Δt* (set to 1 ms) and synaptic delay *Δk* (set to 1 step). Spikes are represented as a variable *s*_*i,k*_ ∈ {0, 1} and generated with probability 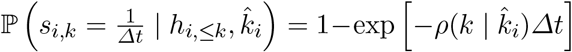 where 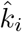 is the time step of the last spike of neuron 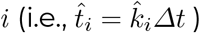 and 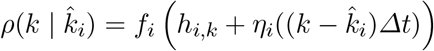 is the firing intensity or hazard.

The history-dependent rate network (Eqs. 3 and 11) can efficiently be simulated forward in time, for a single stimulus presentation, and follows the same time discretization as the spiking network. Specifically, if neuron *i* has a finite refractory kernel *η*_*i*_ of length *k*^max^, such that *η*(*kΔt*) = 0 for all *k* ≥ *k*^max^, the history-dependent firing rate of rate unit *i* in step *k* can be calculated analytically through the expected value of *s*_*i,k*_, conditioned on the rate-based input potential 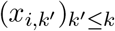. Dropping the *I* notation for readability, we get the per-unit equation

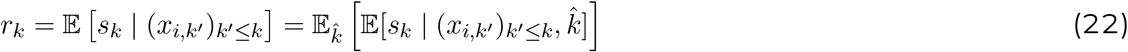

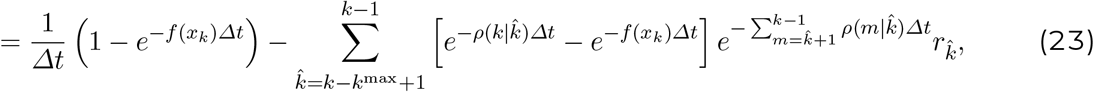

similar to the approach presented in (Herrmann and Gerstner, 2001). This is implemented efficiently using a neuron-specific dynamic buffer 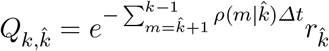 for 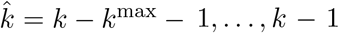, similar to the discrete-time implementation in (Gerstner et al., 2014, Chapter 14.1).

The same approach is used for LIF neurons, with the difference that one needs to keep an additional per-neuron buffer of size at most *k*^max^ − 1 for the history of *x*, also affecting 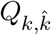.

### Theoretical ISI distribution

From the efficient discrete forward simulation of a history-dependent rate unit *i* it is straight-forward to extract the expected interspike-interval distribution of the spiking neuron *i* it replaces, with negligible additional computational cost for intervals up to the size of the refractory kernel.

Dropping the neuron index *i* notation for readability, the expected neuron-specific ISI distribution is given by

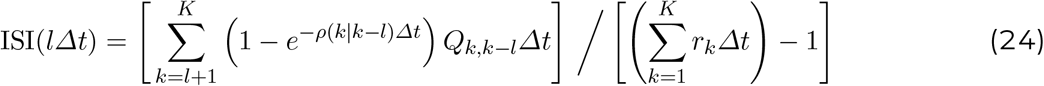

for interval steps *l* = 1, …, *k*^max^ − 1 and with *K* the final time step of the simulation. For a rate unit *i* receiving the same input potential as a spiking neuron, the difference between the spike-based and theoretical ISI distribution vanishes as *K* → ∞.

### Derivation and numerical implementation of the f-I curve for SRM0 and LIF

The f-I curve of neuron *i* refers to the function 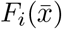 giving the stationary firing rate of neuron *i* as a function of its constant input 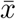. For renewal-type neurons, such as SRM_0_ and LIF, the f-I curve can be deduced from the fact that 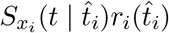 is a probability distribution over the last spike time 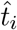. This fact imposes the normalization (Gerstner, 2000)

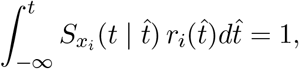

from which we can deduce that for a constant input 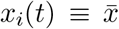, the steady-state firing rate 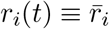 is equal to 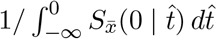. Hence, the f-I curve of the neuron is given by

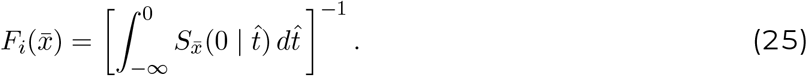

In discrete time, for both SRM_0_ and LIF neurons, the steady-state firing rate for a constant input potential 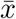 becomes

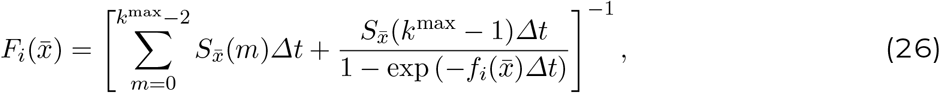

where 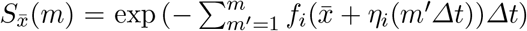 is the survivor function in the stationary state for SRM_0_ neuron *i*, with 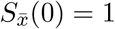 by definition. For LIF, the *η*_*i*_(*m*^*′*^*Δt*) in the survivor function is replaced by 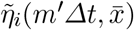 but is otherwise identical.

The function for the discrete f-I curve is used in two ways. First, to initialize the rates of the history-dependent rate network at rest (i.e., *r*_*i*_(*t*) = *F*_*i*_(0) for all *t <* 0). Second, to use in the linear-nonlinear rate networks detailed below.

### Mappings to linear-nonlinear rate networks

We compare three linear-nonlinear rate mappings. In each case, a rate unit *i* is described by a firing rate function *G*_*i*_ that takes the form of a linear-nonlinear cascade (see Eq. 4). The three mappings correspond to three different choices of how the linear filters, or “kernels”, *κ*_*i*_ are obtained from the spiking network parameters, as summarized in Table 2. The non-linearity of each unit *i* is fixed to the f-I curve of spiking neuron *i* (Eq. 25 or Eq. 26).

**Table 2.**
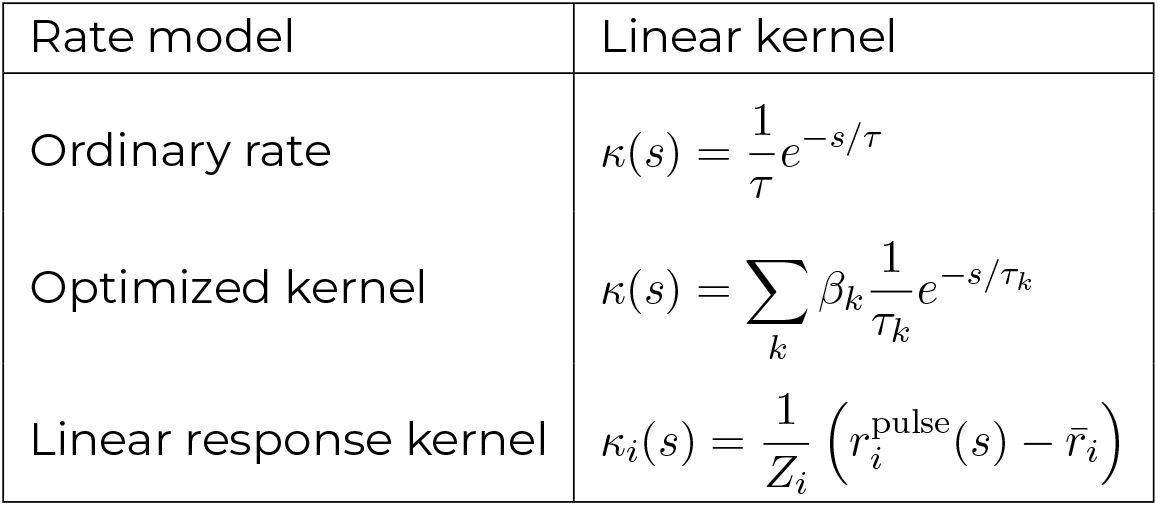
Overview of compared linear-nonlinear rate models, conform Eq. (4).

| Rate model | Linear kernel |
|---|---|
| Ordinary rate | $\kappa(s) = \frac{1}{\tau}e^{-s/\tau}$ |
| Optimized kernel | $\kappa(s) = \sum_{k} \beta_k \frac{1}{\tau_k}e^{-s/\tau_k}$ |
| Linear response kernel | $\kappa_i(s) = \frac{1}{Z_i}\left(r_i^{\mathrm{pulse}}(s) - \bar{r}_i\right)$ |

First, the ordinary rate network (Eq. 5) is obtained by setting all *κ*_*i*_’s to a single exponential function with decay constant equal to the membrane time constant *τ*.

Second, the optimized kernel rate network uses a single parametric kernel that is shared across all units, and is optimized to best reproduce the history-dependent rates. The kernel is represented in an exponential basis with time constants {*τ*_1_, …, *τ*_*K*_} = {2, 4, …, 20} ms.

For kernel optimization, we generate target input-output relations by simulating a reference history-dependent rate network (*N* = 3 × 8000, *P* = 20, either SRM_0_ or LIF) receiving white Gaussian inputs for 10 000 ms. For each unit *i*, we store the rate-based input 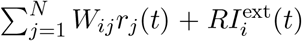 and the resulting output rate *r*_*i*_(*t*). This is used to construct a target potential *h*_target,*i*_(*t*), i.e., the target after linear filtering (*κ*_*i*_ * *I*_*i*_)(*t*), by applying for each time point the inverse of *F*_*i*_ to the history-dependent rate *r*_*i*_(*t*). Kernel coefficients *β*_*k*_ are found through least-squares regression between (*κ* * *I*_*i*_)(*t*) and *h*_target,*i*_(*t*) across all neurons and time points simultaneously. A per-neuron variant was also evaluated, in which each unit *i* has its own kernel coefficients *β*_*i,k*_ found through least-squares regression per unit. In that case, optimization of a given network configuration (some *N* and *P*) was done based on the corresponding history-dependent rate network (same *N* and *P*) receiving white Gaussian inputs for 5 000 ms.

Third, the linear response rate network uses for each rate unit *i* a normalized linear response kernel of the corresponding history-dependent firing rate, linearized around a reference input *I*_0_ (Ostojic and Brunel, 2011). While analytical expressions for the kernel exist for SRM_0_ and LIF (Gerstner, 2000), we used our history-dependent rate implementation to get an efficient approximation by applying a pulse current *I*_*i*_(*t*) = *I*_0,*i*_ + *δ*(*t*)*Δx* of amplitude *Δx* = 0.1 mV at time *t* = 0 relative to a constant input *I*_0,*i*_ to obtain the time course 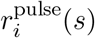, and subtracting the initial condition 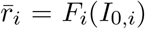. The kernel is normalized by 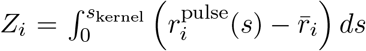. For generalization purposes, the neuron-specific reference input is controlled by a single shared parameter *A*, according to *I*_0,*i*_ = *Aθ*_*i*_ such that *A* = 0 and *A* = 1 yield linear responses around the resting potential and the threshold *θ*_*i*_ (relative to rest), respectively. The parameter *A* was optimized through grid search on a network of size *N* = 3 × 8000 driven by *P* = 20 white Gaussian input signals for *T* = 500 ms. *A* was systematically varied between −1 and 2 in steps of 0.1, with the goal to minimize the RMSE between the output of a reference spiking network and a corresponding linear response rate network, which yielded *A* = 1.6 and *A* = 0.2 for SRM_0_ and LIF, respectively.

For each rate network, corresponding input potentials *x*_*i*_(*t*) are obtained through Eq. (3). With the exception of the ordinary rate network, input potentials *x*_*i*_(*t*) do not explicitly enter in the linear-nonlinear rate function (Eq. 4) and are computed solely for the purpose of comparison with the spike-based input potentials *h*_*i*_(*t*). Numerical simulations make use of the discrete f-I function (Eq. 26), which for computational efficiency are sampled at regularly spaced intervals, from 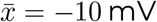 to 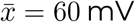 (relative to rest) in steps of 0.5 mV, and linearly interpolated at runtime. Values outside this range are clipped to the nearest point.

### Sparse networks and weight similarity

Sparse networks are constructed with a fixed number of incoming connections per neuron *K* ≤ *N*. Specifically, we take a structured weight matrix as before but with normalization 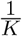 instead of 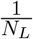, from which for each neuron (i.e., for each row in *W* ^(*l*)^) exactly *N*_*L*_ *K* randomly selected connections are pruned.

Shared variability arises from pairs of postsynaptic neurons sampling from the same presynaptic neurons. We quantify this using the cosine similarity in incoming weight vectors between different pairs of neurons.

## Supplementary Information

### Supplementary Note 1: Concentration argument

In this Supplementary Note, we present a concentration argument explaining how the history-dependent rate model can capture the dynamics of heterogeneous spiking networks, thereby providing a justification for the virtual-population trick we used in the derivation of the model. The concentration bound presented in the following extends a previous result for networks linear-nonlinear Poisson spiking neurons (Schmutz et al., 2025) to networks of renewal-type spiking neurons.

To demonstrate the spike-to-rate concentration phenomenon, we focus on the case of a single-layer feedforward network consisting of *N* SRM_0_ neurons sending spikes to a single output neuron. This setup is similar to that studied in Fig. 2. The input potentials of the output neurons evolves according to

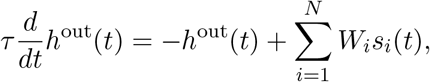

where the *W*_*i*_’s are the synaptic weights from spiking neuron *i* to the output neuron. For any index *i*, the spikes *s*_*i*_(*t*) of input neuron *i* are generated from a SRM_0_ spiking neuron model, i.e.,

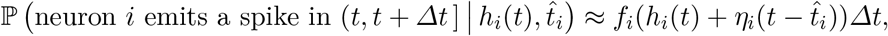

where 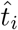 is the last firing time of neuron *i* and *h*_*i*_(*t*) is its input potential (see main text for the full definition of the model). Here, the input potential is arbitrary; it can be for example be given by a linear readout of low-dimensional inputs, as in Fig. 2 The spike-emission noise is assumed to be independent between neurons, conditioned on the input potentials (*h*_1_(*t*), *h*_2_(*t*), …, *h*_*N*_ (*t*)).

We want to compare the spike-based input potential, *h*^out^(*t*), with its rate-based counterpart, *x*^out^(*t*), given by the history-dependent rate model,

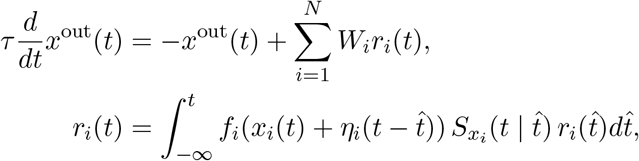

while fixing *x*_*i*_(*t*) = *h*_*i*_(*t*) for all *i* = 1, …, *N*, i.e., the input spiking neurons and the input rate units share the same input potentials. We also assume that the initial condition of the spiking network and the rate model are matched: for all 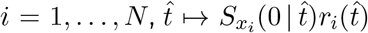 is a probability distribution over (, 0], and the last firing time of neuron *i*, 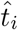, is sampled randomly from this distribution, independently for each neuron; in addition, we assume that *h*^out^(0) = *x*^out^(0). For any given recording time *T >* 0, we quantify the distance between the spiking network and the rate model with the expected mean-squared error

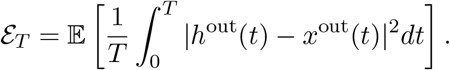

#### Theorem 1

*Let us assume that that soft-threshold function f*_*i*_ *are bounded, and let us write* 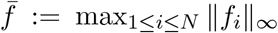. *Furthermore, let us write* **W** := (*W*_1_, *W*_2_, …, *W*_*N*_)^T^ *the vector of input weights. Then*,

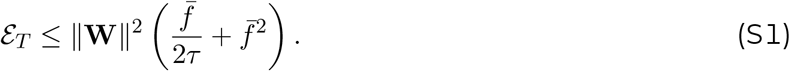

The theorem says the the mean-squared error depends on the squared Euclidean norm of the input weight vector. Importantly, the bound Eq. (S1) does not require the neurons to be identical; spike-emission noise simply has to be conditionally independent between neurons. Here are some additional remarks:

1. The bound Eq. (S1) is a non-asymptotic bound in the sense that is holds for networks of any finite size *N*.
2. For the error *ℰ*_*T*_ to tend to zero as *N→ ∞*, it suffices that the norm of the input weights ∥**W**∥ tends to zero as *N→ ∞*. This is strictly weaker than requiring a 1*/N* scaling of the weights. For example, with the scaling *P* = *N* ^1*/*3^ used in Fig. 2, individual weights scale as 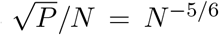 (which is strictly slower than 1/*N*), and this is enough for the norm ∥**W**∥ to tend to zero as *N* (at rate 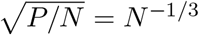). Hence, the bound Eq. (S1) is not a mere law-of-large-number-type result but a more general, concentration of measure result.
3. While the result is presented here for the case of SRM_0_ neurons, the same types of arguments apply for networks of LIF neurons with escape noise.

*Proof*. Solving the differential equation for *h*^out^(*t*) and *x*^out^(*t*), we obtain

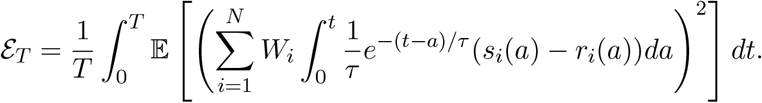

Developing the square,

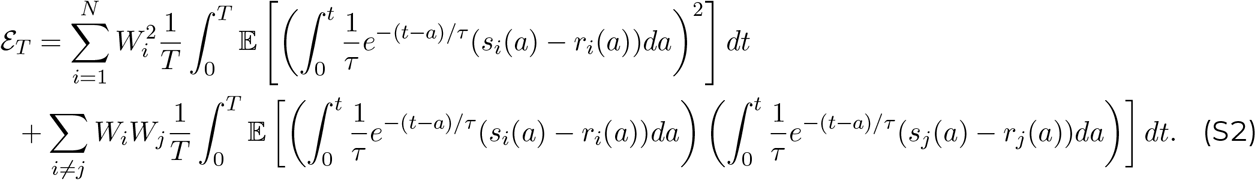

For the cross terms *i ≠ j*, by the conditional independence of the spike-emission noises,

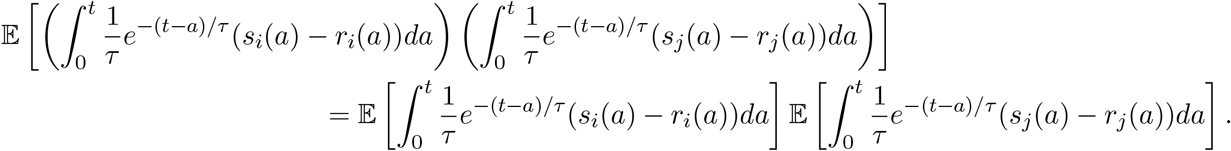

We recall that *r*_*i*_(*t*) is, by definition, the expected time-varying firing rate of neuron *i* conditioned on its input potential. Hence,

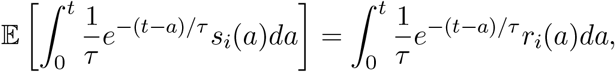

which implies that the cross terms on the right-hand side of Eq. (S2) are zeros, and we obtain

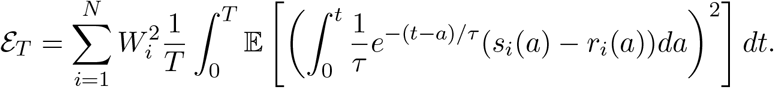

To bound the variance term on the right-hand side above, we first use the fact that

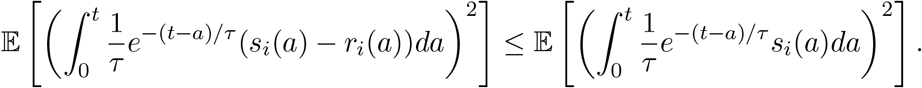

Let 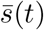 be a spike train generated by a homogeneous Poisson process with rate (intensity) 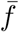. Since 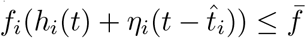,

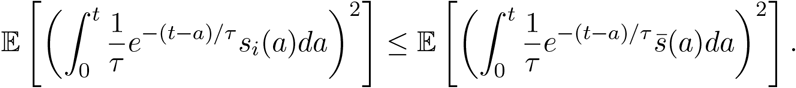

By the standard theory of Poisson processes,

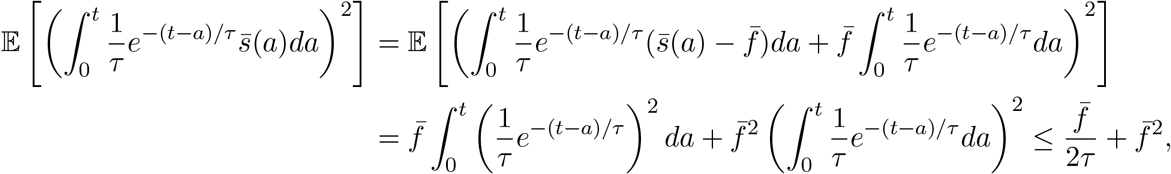

which yields the desired bound.

---

**Figure S1.**
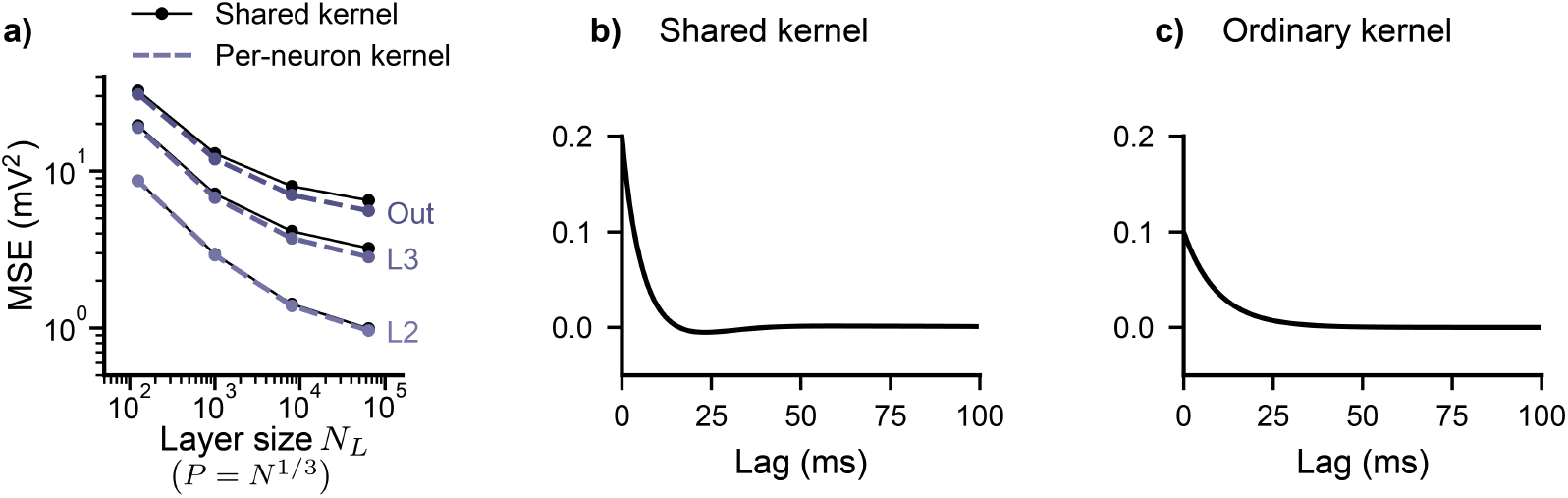
Shared optimized kernel mapping is comparable to per-neuron optimized kernel mapping. **a)** Mismatch compared to SRM_0_ spiking network across layers (L2, L3, Out) measured in MSE (mean across neurons/outputs and *n* = 6 initializations) for varying number of neurons per layer *N*_*L*_ on a log-log scale, for two linear-nonlinear mappings using kernel fitting. **b)** The shared kernel found through parametric optimization for SRM_0_ neurons, see Methods. **c**) The ordinary kernel used to construct the ordinary rate network, shown for reference.

**Figure S2.**
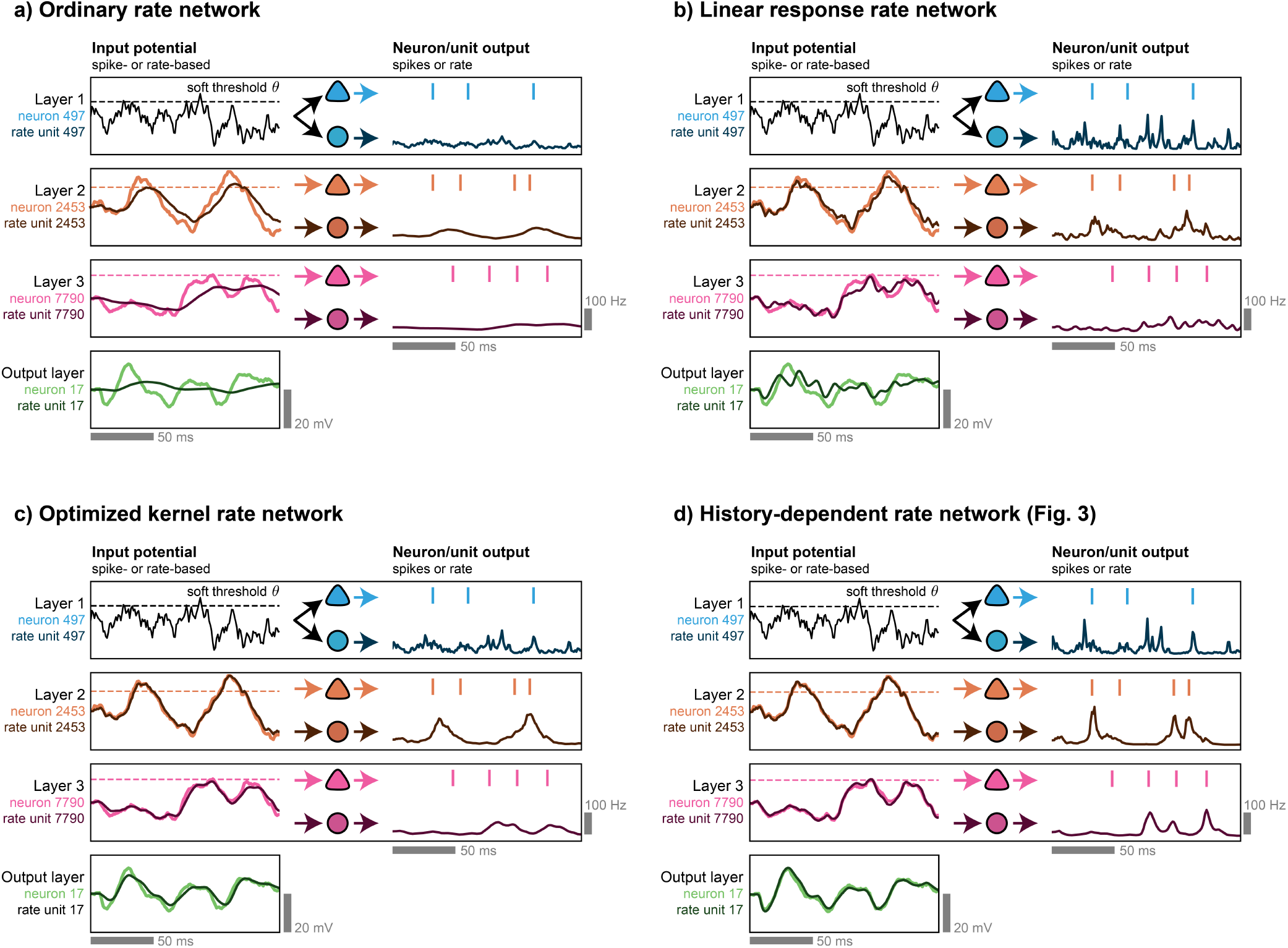
Dynamics of compared rate networks. Dynamics of all four evaluated rate networks in describing the dynamics of the spiking network of Figure 3. Each rate network forms its own closed system with rate-based interactions. Data for history-dependent rate network (copied from Fig. 3b) included for comparison. See Figure 3 for details.

## Notes

### Competing Interest Statement

The authors have declared no competing interest.

